# Temporal Analysis of T-Cell Receptor-Imposed Forces via Quantitative Single Molecule FRET Measurements

**DOI:** 10.1101/2020.04.03.024299

**Authors:** Janett Göhring, Florian Kellner, Lukas Schrangl, René Platzer, Enrico Klotzsch, Hannes Stockinger, Johannes B. Huppa, Gerhard J. Schütz

## Abstract

Mechanical forces acting on ligand-engaged T-cell receptors (TCRs) have previously been implicated in T-cell antigen recognition, yet their magnitude, spread, and temporal behavior are still poorly defined. We here report a FRET-based sensor equipped with a TCR-reactive single chain antibody fragment, which was tethered to planar supported lipid bilayers (SLBs) and informs most directly on the magnitude and kinetics of TCR-imposed forces at the single molecule level. When confronting T-cells with gel-phase SLBs we observed both prior and upon T-cell activation a single, well-resolvable force-peak of approximately 5 pN and force loading rates on the TCR of 1.5 pN per second. When facing fluid SLBs instead, T-cells still exerted tensile forces yet with threefold reduced magnitude and only prior to but not upon activation. Our findings do not only provide first truly molecular information on TCR-imposed forces within the immunological synapse, they also recalibrate their significance in antigen recognition.

## INTRODUCTION

A remarkable feature of the vertebrates’ immune system is its inherent ability to distinguish harmful from harmless based on the primary antigen structure. T-cells embody this trait of adaptive immunity through their unique detection of antigens, which are displayed as peptide fragments in the context of the products of the major histocompatibility complex (MHC) on the surface of antigen-presenting cells (APCs). The formation of a productive immunological synapse, the area of contact between a T-cell and an APC (1), is driven by T-cell antigen receptors (TCRs) on the T-cell binding to peptide/MHC complexes (pMHC) on the APC. Importantly, T-cells specifically detect the presence of even a single antigenic pMHC molecule among millions of structurally similar yet non-stimulatory pMHCs (2). This is even though nominal pMHC-TCR interactions are of rather moderate affinity, at least when measured *in vitro*. Despite considerable interest, the molecular, biophysical and cellular mechanisms underlying this phenomenal sensitivity and specificity are not well understood.

Tensile forces have been implicated in the discrimination of antigenic peptides from non-activating pMHCs, especially if T-cells gauge TCR-proximal signaling based on TCR-pMHC bond lifetimes as is proposed by the kinetic proofreading model (3–6). Over the course of the last decade such forces have garnered significant attention and have even become regarded an instrumental trigger in T-cell antigen detection (7, 8). With the use of a single-molecule Förster Resonance Energy Transfer (FRET)-based live cell imaging assay, we have observed synaptic unbinding between TCR and pMHC with significantly increased off-rates compared to TCR-pMHC binding in solution (9), possibly as the result of cellular motility. Additionally, cell mechanical (10, 11) and *in vitro* (12) studies have revealed altered TCR-pMHC unbinding under force, yielding indications for the relevance of both catch bonds (10) as well as slip bonds (12) for the antigen discrimination process.

When confronted with beads that had been functionalized with TCR-ligands, T-cells were observed to undergo calcium signaling upon mechanical bead retraction (7, 13). Activation thresholds were found markedly reduced when pMHC-coated beads were moved tangentially to the T-cell surface (14). In line with this, T-cells were reported to push and pull against model APCs in the course of antigen-driven activation (15, 16). It has also been proposed that T-cells actively probe the elastic properties of the substrate (17), and global shear forces of up to 150 pN were observed when T-cells had been activated via antigen-functionalized elastomeric pillar arrays (18) or polymers with embedded beads (19).

These and related findings prompted efforts to quantify mechanical forces directly within the immunological synapse. Experiments involving the use of DNA-based tension sensors have given rise to the suggestion that forces larger than 12 pN were acting on ligand-engaged TCRs when T-cells were confronted with antigenic glass-immobilized pMHCs (20). Forces amounting to up to 4.7 pN were reported for T-cells that interact with planar glass-supported lipid bilayers (SLBs) featuring pMHCs anchored to gold particles (21). Furthermore, when an adjustable upper force limit was imposed on TCR-pMHC bonds with the use of rupturable DNA tension gauge, the recruitment of the TCR-proximal tyrosine kinase ZAP70 to the plasma membrane, a process that is integral to early T-cell signaling, correlated well with the maximal amplitude of the allowed force (20).

However, current ensemble measurements render it challenging to assess pulling forces acting on individual TCRs. Typically, multiple bonds are simultaneously subjected to strain, and it is not clear whether the DNA-based tension sensors are arranged in parallel or zipper-like configuration (22). Furthermore, when considering the mechanism underlying the function of such sensors, there is a strong dependence of the unzipping force on the loading rate (23) as well as on the duration of the applied force (24), which constitute *a priori* unknown parameters. If given sufficient time, all DNA duplexes will eventually separate when any force is applied. Last but not least, ensemble-based readout modalities fail to afford kinetic force profiles acting on individual receptor-ligand bonds which hence remain undetermined.

In this study, we report the development and application of a quantitative FRET-based force sensor for application within the immunological synapse, which operates at the single-molecule level. For a spring element, we employed a peptide derived from the spider silk protein flagelliform comprised of 25 amino acids with known elastic properties (25, 26). The spring peptide was anchored to either gel- or fluid-phase SLBs and conjugated either to a single chain antibody fragment (scF_V_) derived from the TCRβ-reactive H57 monoclonal antibody (H57-scF_V_) to serve as a high-affinity artificial TCR-ligand, or to MHC loaded with a nominal peptide as the natural TCR-ligand. In its collapsed configuration the sensor spans a distance of less than 8nm and is therefore compatible with length restrictions of approximately 13 nm as they apply within the immunological synapse (27, 28). When engaging TCRs with the high-affinity ligand H57-scF_V_, we observed 5 to 8 pN average forces per TCR for T-cells contacting gel-phase SLBs. Compared to H57-scF_V_, force readings obtained with the use of the natural ligand pMHC showed a broader distribution shifted towards lower forces amounting to about 2 pN. Of note, only barely measurable TCR-imposed forces below 2 pN were detected when T-cells interacted with fluid-phase SLBs. Since both gel-and fluid-phase SLBs support T-cell stimulation, our measurements imply that perpendicular tensile forces smaller than 2 pN are already effective during the initiation of TCR signaling. Our findings render it furthermore highly unlikely that forces, at least to an extent they are measurable with our FRET-based system, are required for continual TCR-signaling at later stages of synapse formation.

## RESULTS

### Design of the force sensor

We designed the molecular force sensor (MFS) in a fashion that rendered it compatible with the use of protein-functionalized SLBs. The latter has been widely employed as surrogates of antigen presenting cells (1), especially with the incorporation of nickel-chelating lipids, such as 1,2-dioleoyl-sn-glycero-3-[(N-(5-amino-1-carboxypentyl)iminodiacetic acid)succinyl] (Ni^2+^) (Ni-NTA-DGS), which supports decoration of SLBs with poly-histidine (His)-tagged proteins of choice in experimentally definable densities (9). The use of SLBs also allows for the application of Total Internal Reflection-(TIR) based imaging modalities, which give rise to sufficiently low background noise for recording single molecule traces.

Monovalent recombinant streptavidin (mSAv) was utilized in our study as anchor unit for the force sensor to be embedded in the SLB (Fig. 1A, B). To ensure stoichiometric anchorage we engineered the tetrameric mSAv complex to contain one “alive”, i.e. biotin-binding subunit and three “dead” subunits, which were mutated to no longer bind biotin. Instead, dead subunits featured a tag comprised of six histidine residues each (mSAv-3xHis_6_, for more details refer to Methods section) for attachment to SLBs employed in our study, which contained 2% Ni-NTA-DGS for functionalization with poly-histidine-tagged proteins. Depending on the experiment, the carrier lipids (98%) were either 1,2-dipalmitoyl-sn-glycero-3-phosphocholine (DPPC) giving rise to immobile, gel-like SLBs which allowed for shear and pulling forces to take effect, or 1-palmitoyl-2-oleoyl-glycero-3-phosphocholine (POPC), which enabled high lateral protein mobility, and were as a consequence permissive for pulling forces only.

**Figure 1:**
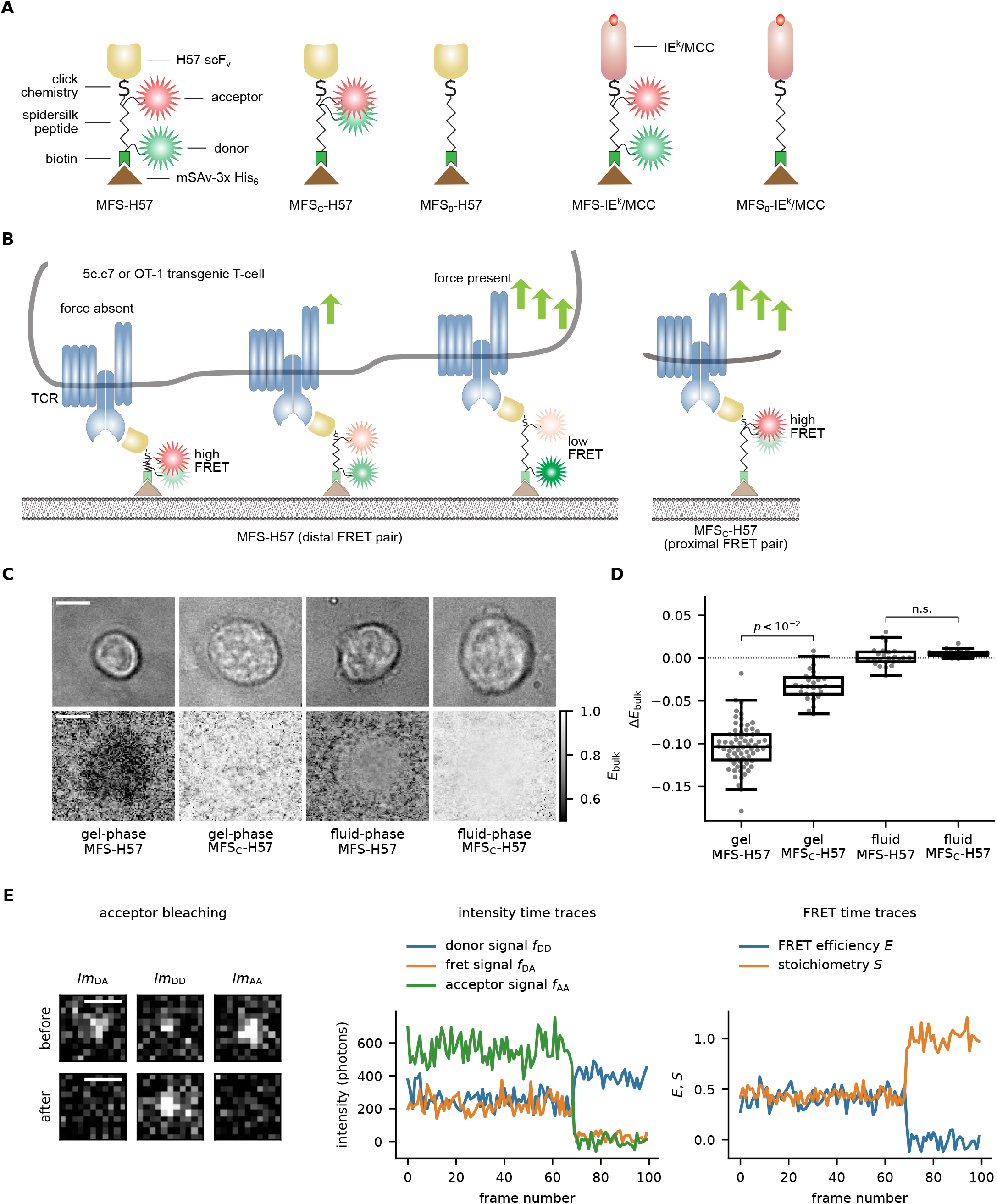
FRET-based sensor designed to detect forces on a single-molecule level. (**A**) Schematic illustration of the force sensor constructs MFS-H57, and MFS-IE^k^/MCC, the control construct MFS_C_-H57, and the unlabeled constructs MFS_0_-H57 and MFS_0_-IE^k^/MCC. (**B**) Exertion of forces stretches the spring core module, which reduces FRET yields for the molecular force sensor (here: MFS-H57) but not for the control force sensor (here: MFS_C_-H57). (**C**) Representative images illustrating FRET efficiencies determined on a pixel-by-pixel basis via DRAAP on gel-phase (DPPC) and fluid-phase (POPC) SLBs with the use of the MFS or the MFS_C_ construct (bottom row), and the corresponding transmission light microcopy images (top row). (**D**) Statistical analysis of ensemble FRET measurements. Each data point corresponds to a single DRAAP experiment of up to three cells in the field of view; boxes indicate the interquartile range (first quartile to third quartile), whiskers extend 1.5 times the interquartile range from the first and third quartile, the line indicates the median. A significant reduction in FRET efficiency (*ΔE_bulk_*) was only observed for MFS on gel-phase SLBs (n.s., not significant; two-sample Mann Whitney U test). n = 68 cells for MFS on gel-phase SLBs, n = 24 for MFS on fluid-phase SLBs, n = 27 for MFS_C_ on gel-phase SLBs, n = 21 for MFS_C_ on fluid-phase SLBs. (**E**) Representative single-molecule signals and time traces. The left panel depicts the fluorescence signals prior to and after photobleaching of the FRET acceptor. The middle panel shows single-molecule brightness values of the FRET donor, the FRET signal itself and the FRET acceptor. Bleaching of the acceptor could be observed as a correlated step-wise decrease of FRET and FRET acceptor intensities and an increase in FRET donor intensity. The right panel shows the stoichiometry and FRET efficiency. Scale bars: 5 µm (panel C), 1 µm (panel E).

We designed the MFS to support direct visualization and quantification of single-digit piconewton forces via FRET with a temporal resolution in the millisecond range, as they were expected to occur between receptor-ligand pairs within the immunological synapse (Fig. 1A). Integral to the MFS were two corresponding fluorophores acting as a FRET pair and flanking the sensor’s core which were derived from the spider silk protein flagelliform and consisted of five repeats of the peptide motif GPGGA (25, 29). We expected the spring to collapse with high FRET yields associated with the absence of force, yet upon force application to give rise to reduced FRET yields due to an increased distance separating the two fluorophores (26).

For a quantitative readout, the core module of the MFS had to (i) be amenable to site-specific modification with fluorophores of choice, (ii) afford directional conjugation of the spring module to the T-cell ligand and (iii) provide anchorage to SLBs (Fig. 1A, B). To fulfill these needs, we synthesized the spring peptide with an unpaired cysteine and a single lysine residue for fluorophore conjugation via maleimide and succinimidyl chemistry, respectively. The peptide’s N-terminus was biotinylated to omit the N-terminal amino group and to confer binding of the peptide to SLB-resident mSAv-3xHis_6_ (30). C-terminal incorporation of an ε-N_3_ modified lysine residue in the sensor’s core facilitated covalent attachment of site-specifically dibenzylcyclooctyne-(DBCO-) conjugated TCR-ligand via biorthogonal copper-free click chemistry (31).

As a TCR-engager we generated two constructs: first, we utilized the TCRβ-reactive recombinant H57-scF_V_ (MFS-H57; for construction see Supplementary Fig. 1A-D), as we anticipated this to give rise to stable TCR-bonds, which unlike pMHC-TCR interactions sustain the tensile forces applied in the synapse without rupture. It is known that monomeric membrane-bound H57 activates T-cells with similar activation thresholds as agonistic pMHC (32, 33). Second, we also functionalized the force sensor with MHC class II molecule IE^k^ loaded with a moth cytochrome c (MCC, IE^k^/MCC) peptide (MFS-IE^k^/MCC; for construction see Supplementary Fig. 1A-D), the nominal and physiological antigen of 5c.c7 TCR-transgenic CD4+ T-cells, which we employed predominantly in this study (see below). Importantly, the N-terminal biotin residue within the MFS core allowed for stoichiometric coupling to mSAv-3xHis_6_ present on the SLB (Supplementary Fig. 1E).

In some experimental settings we aimed to measure forces exerted by individual TCRs under moderate levels of antigenic stimuli (see below). To this end we diluted the fluorophore signals with an unlabeled force sensor termed MFS_0_-H57 or MFS_0_-IE^k^/MCC. In addition, a stretch-insensitive version of the sensor was constructed to serve as a control (MFS_C_-H57). Here the FRET acceptor and FRET donor fluorophores were separated by only a single amino acid (Fig. 1A, B).

Integrity of all employed constructs was confirmed by SDS-PAGE (Supplementary Fig. 1E). When anchored to SLBs all constructs gave rise to robust T-cell activation as monitored via the rise in intracellular calcium (Supplementary Fig. 1F-G).

### Forces in the immunological synapse visualized at the ensemble level

CD4^+^ helper T-cells were isolated prior to their experimental use from the spleen and lymph nodes of 5c.c7 TCR transgenic mice and stimulated *ex vivo* for one week with antigenic peptide to promote differentiation into antigen-experienced T-cell blasts. These were then seeded onto SLBs functionalized with Intercellular Adhesion Molecule 1 (ICAM-1), B7-1, and MFS-H57 or MFS_C_-H57 (50-100 molecules per µm^2^). We first confronted T-cells with gel-phase SLBs prepared from DPPC, with associated proteins showing minimal diffusion (*D* ≤ 1.5 x 10^-4^ µm^2^ s^-1^; Supplementary Fig. 2A). As shown by calcium imaging (Supplementary Fig. 1F and Supplementary Fig. 3A) and the analysis of immune synapse recruitment of ZAP70 (Supplementary Fig. 3D), T-cells responded to such SLBs in a highly sensitive manner. Peak calcium signals were identical with that of the positive control, which involved stimulation via SLB-anchored IE^k^/MCC, and elevated calcium flux was sustained for at least 15 minutes. We noticed a marginally delayed onset of T-cell activation triggered by force sensor-constructs when compared to that of the positive control, possibly reflecting slightly altered stimulation conditions within the synaptic space constraints (34). Furthermore, the high viscosity inherent to DPPC SLBs prevented clustering of TCR-engaged SLB-resident MFS-H57 (Supplementary Fig. 2B), which we have previously observed to affect antibody-mediated T-cell activation (32).

Ensemble FRET efficiencies (*E_bulk_*) were quantified via Donor Recovery After Acceptor Photobleaching (DRAAP) by comparing the average donor signal in a chosen region of interest before and after the photoablation of the acceptor fluorophore. A substantial reduction in *E_bulk_* underneath the T-cell indicated the elongation of the force sensor (Fig. 1C) which we did not observe when applying the control construct. For the statistical analysis, we calculated the difference *ΔE_bulk_* by subtracting the region-matched E_bulk_ recorded on a functionalized SLB in the absence of T-cells from E_bulk_ underneath the T-cell, yielding a reduction *ΔE_bulk_* of −0.104 (median). In contrast, the control construct showed a significantly smaller FRET-reduction *ΔE_bulk_* of −0.033 (Fig. 1D).

To account for the comparably high mobility of MHC, ICAM-1 and B7-1 present on the surface of living APCs (35), we confronted T-cells with fluid-phase SLBs which were prepared from 1-palmitoyl-2-oleoyl-glycero-3-phosphocholine (POPC) and subsequently functionalized with proteins in the same fashion as the gel-phase SLBs. SLB-anchored MFS-H57 that underwent lateral diffusion with *D* equaling 0.72 ± 0.02 µm^2^ s^-1^ (Supplementary Fig. 2A), were rapidly recruited into TCR-microclusters, transported to the center of the synapse (Supplementary Fig. 2B) and resulted in a robust calcium response and ZAP70 recruitment (Supplementary Fig. 1F and Supplementary Fig. 3B, D), albeit with a slightly delayed calcium flux in case of gel-phase SLBs. However, we did not observe significant reductions in FRET within synaptic MFS-H57 (*ΔE_bulk_*=0.000 median for the MFS-H57 and *ΔE_bulk_*=0.005 for the MFS_C_-H57) (Fig. 1D).

Together, ensemble-based FRET experiments indicate the presence of tensile forces exerted by T-cells via their TCRs. FRET-changes were, however, only detectable when T-cells interacted with gel-but not fluid-phase membranes. Our observations suggest that TCR-exerted force fields acted foremost tangentially to the T-cell surface: in such cases, gel-phase membranes provided adequate counterforces to stretch the MFS-H57, while fluid-phase membranes would allow for MFS-H57 dragging without any FRET changes.

### Forces in the immunological synapse at the single molecule level

While informative on a qualitative level, ensemble FRET measurements via the MFS are only of limited use for quantitative analysis for a number of reasons. First and foremost, any information regarding the kinetics and amplitude of single-molecular forces is masked in the ensemble average, which includes in addition to sensors under tension also contributions from unbound force sensors or sensors bound to TCR-enriched microvesicles (36), rendering the quantification of individual TCR-exerted forces impossible. Moreover, defective sensors may contribute to observed ensemble FRET efficiencies in ways that are difficult to account for or predict. Last, but not least, molecular fluorophore enrichment within the synapse in the course of T-cell activation could affect the FRET-readout.

To circumvent these limitations altogether, we carried out single molecule microscopy experiments, which allowed us to filter out defective sensors and quantify forces between individual TCRs and SLB-anchored MFS-H57 in time and space with the use of both gel- and fluid-phase SLBs for T-cell stimulation. For this, T-cells were seeded onto SLBs functionalized with ICAM-1, B7-1, the unlabeled sensor MFS_0_-H57 (50-100 µm^-2^), and low concentrations of the MFS-H57 or MFS_C_-H57 (<0.01 µm^-2^). As shown in Supplementary Fig. 3, the stimulatory potency of such SLBs was verified by monitoring the calcium response and ZAP70-recruitment of interacting T-cells.

After low enough MFS-H57 densities were chosen to allow for the emergence of well isolated single molecule signals within the immunological synapse, we imaged the donor and acceptor channel at alternating excitation over time and calculated the FRET efficiency *E* via sensitized emission, taking into account the corresponding cross-talk signals (see Materials & Methods for a detailed description, and Fig. 1E for exemplary traces) (37). We also determined the stoichiometry parameter *S*, which measures the fraction of donor molecules in donor-acceptor complexes (37). When plotting *S* against *E* for each observed data point, we found a single population centered around *S∼0.5*, testifying to the robustness of the filtering steps for single and functionally intact MFS-H57 (Supplementary Fig. 4; see Supplementary Figure 5 for more details on the procedure). In the absence of T-cells we observed for MFS-H57 and MFS_C_-H57 pronounced maxima at *E* = 0.87 and *E* = 1.00, respectively, which represented the FRET efficiencies of collapsed springs. The higher FRET efficiency recorded for the MFS_C_-H57 construct resulted in all likelihood from the shorter distance within the FRET donor-FRET acceptor dye pair, regardless of the spring extension.

Inspection of the immunological synapse revealed considerable differences in the spectral behavior of the two MFS constructs: while MFS_C_-H57 showed FRET efficiencies that were almost identical to those recorded in the absence of cells (E = 0.99), for MFS-H57 we detected a second peak at *E* = 0.41 with a weight of 21% in addition to the high-FRET peak at *E* = 0.87. This low-FRET peak corresponded to MFS-H57 subjected to force-induced elongation via ligated TCRs.

To read out the applied tensile forces from determined FRET efficiencies we factored in the elastic properties of the flagelliform force sensor, which were previously determined in combined single molecule force spectroscopy and FRET experiments on a similar construct (26). We inferred the relationship between FRET efficiency and the tensile force acting on the sensor used in our study (see Materials and Methods for a detailed description of the calibration) and calculated an average pulling force *F* of 4.9 ± 0.3 pN for the low FRET peak at *E* equaling 0.42 ± 0.03 (Fig. 2A, first panel). Expectedly, pulling forces were abrogated when inhibiting actin polymerization by cytochalasin D (Supplementary Fig. 6A).

**Figure 2:**
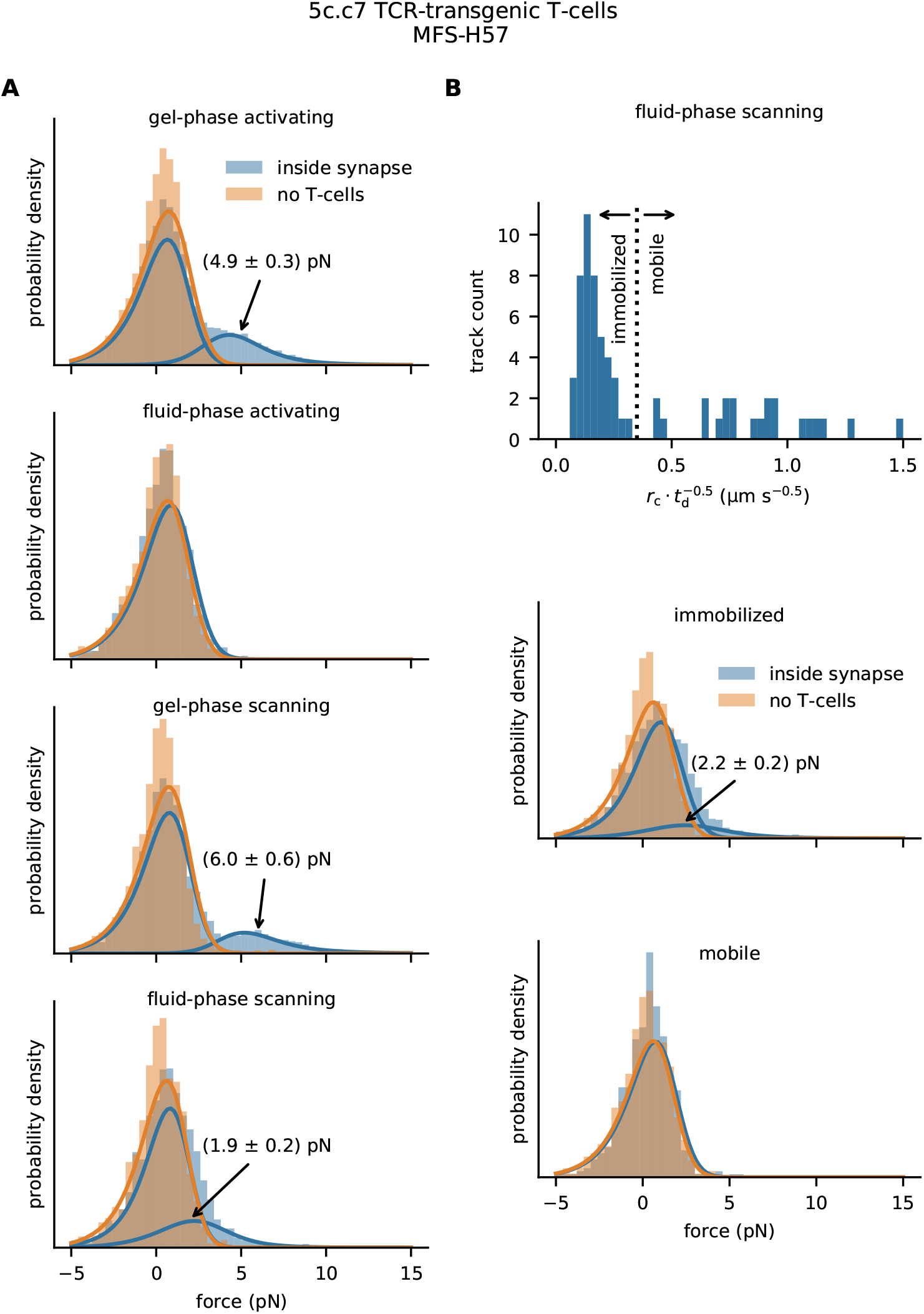
Single-molecule pulling forces exerted via individual TCRs on the H57-functionalized molecular force sensor. (**A**) Single-molecule TCR-imposed forces determined within synapses of activated or scanning T-cells confronted with gel-phase (DPPC) or fluid-phase (POPC) SLBs. Experiments were performed on 5c.c7 T-cells using sensors functionalized with H57. A high force peak emerged for activated or scanning T-cells confronted with gel-phase SLBs at *F* = 4.9 ± 0.3 pN and *F* = 6.0 ± 0.6 pN, which corresponded to 21% and 15% of observed events, respectively. For T-cells scanning fluid-phase SLBs we observed an additional high force peak at *F* = 1.9 ± 0.2 pN with 25% weight. n = 15556 (593 movies, each containing ≥ 1 cell) for MFS on gel-phase SLBs in contact with activated T-cells, n = 12657 (195 movies) for MFS on fully stimulatory gel-phase without T-cells, n = 6006 (453 movies) for MFS on gel-phase SLBs including scanning T-cells, n = 5502 (165 movies) for MFS on gel-phase SLBs (non-stimulatory) without T-cells, n = 1161 (293 movies) for MFS on fluid-phase interacting with activated T-cells, n = 1528 (90 movies) for MFS on stimulatory fluid-phase without T-cells, n = 2147 (333 movies) for MFS on fluid-phase SLBs in contact with scanning T-cells, n = 1796 (70 movies) for MFS on non-stimulatory fluid-phase without T-cells. (**B**) Bound MFS were distinguished from unbound MFS on fluid-phase SLBs under scanning conditions by means of the single molecule mobility as described in Supplementary Fig. 8. The dashed line indicates the chosen boundary drawn between mobile and immobilized MFS. Unbound MFS (termed “mobile”) gave rise to a force distribution showing no significant difference from the control scenario recorded without T-cells. Bound MFS (termed “immobilized”) featured a high-force shoulder similar to Fig. 4A, fourth panel.

To assess the dynamic range of our sensor, we estimated the minimum and maximum FRET efficiency of the low FRET peak that could be measured with the use of the MFS-H57. The smallest detectable FRET value *E_min_* was set by the donor-acceptor distance at maximum load, the Förster radius of the chosen FRET pair, and the signal to noise ratio of the data. As is laid out in more detail in the Methods section, we estimated *E_min_* to amount to 0.11, which relates to a tensile force *F_max_* of 10 pN. Furthermore, the largest detectable FRET value *E_max_* was limited by the overlap with the high FRET peak at *E* = 0.87. With the observed proportions of the two peaks we estimated *E_max_* to ∼ 0.8, which related to a minimum detectable force *F_min_* of about 0.9 pN. We therefore conclude that the observed force peaks fell well within the sensor’s dynamic range, and that they corresponded to a not fully extended spring element.

In line with the results of the ensemble measurements (see above) we did not observe a high force peak for sensors within synapses when applying fluid-phase SLBs prepared from POPC (Fig. 2A, second panel, Supplementary Fig. 7, and Supplementary Table 1). MFS-H57 binding to the TCR was confirmed in single molecule tracking experiments in which we found the majority of MFS-H57 (58%) to exhibit a substantially reduced mobility underneath T-cells. In control experiments involving human Jurkat T-cells, which do not engage the MFS-H57 for lack of H57-scF_V_-reactivity, we did not observe synaptic MFS-H57 immobilization (Supplementary Fig. 8).

### TCR-imposed forces under scanning conditions

Next, we aimed to assess the degree to which the high number of molecular bonds formed between the SLB and the T-cell disperses cellular forces over multiple TCR-ligand bonds while lowering the forces acting on individual MFS molecules and their associated TCRs. Of note, such parallel arrangement of multiple TCR-ligand bonds may very well apply to activated T-cells featuring a fully established synapse, yet to a much lesser extent to T-cells scanning APC surfaces for antigenic ligands, where isolated TCR-ligand interactions become likely subject to larger tensile forces. To discriminate these scenarios, we seeded T-cells onto ICAM-1- and B7-1-functionalized SLBs featuring MFS-H57 in low densities below the activation threshold (<0.01 molecules per µm^2^) (Supplementary Fig. 3A, B & D). Again, we detected a distinct high force peak corresponding to 6.0 ± 0.6 pN for T-cells confronted with gel-phase SLBs, which slightly exceeded the force peak measured for fully activated T-cells (Fig. 2A, third panel). We noticed a shoulder towards lower applied forces corresponding on average to 1.9 ± 0.2 pN for T-cells contacting fluid-phase SLBs, (Fig. 2A, fourth panel). Next, we took advantage of the higher mobility of unbound sensors when compared to that of bound sensors, and specifically analyzed the force histograms for bound and unbound MFS-H57 (Fig. 2B). As expected, unbound, i.e. mobile MFS-H57 did not register any tensile forces, while immobilized, i.e. TCR-engaged sensors gave rise to a shoulder in the force peak corresponding to 2.2 ± 0.2 pN, a value that was found in agreement with the global distribution of the combined data and about three-fold below that measured for TCRs interacting with immobile MFS-H57. Together, the data acquired at low MFS-H57 concentrations imply that perpendicular pulling forces exerted via the TCR are not a consequence of T-cell activation and may well precede the actual activation step.

### Spatiotemporal analysis of TCR-imposed forces

An interesting aspect of our observations on gel-phase membranes relates to the similarity between tensile forces applied by T-cells scanning for antigenic peptide and those imposed by activated T-cells. To further scrutinize the obtained data, we pooled the data in sliding windows of 5 minutes after initial T-cell-SLB contact. The results revealed remarkable differences between scanning and activated T-cells. For activated T-cells (Supplementary Fig. 9A), we observed the emergence of the high force peak only after ∼10 minutes at 5.6 ± 0.1 pN, where it remained afterwards. Tensile forces occurred both in the cell periphery as well as in central regions of the immunological synapse (Supplementary Fig. 10A). Quite in contrast, scanning T-cells exhibited the high force peak immediately following initial SLB contact (Supplementary Fig. 9B). Interestingly, the average TCR-imposed pulling force declined over time from 7.5 ± 0.2 pN within the first 5 minutes to 6.3 ± 0.2 pN 5 to 10 minutes after contact, and later down to low force magnitudes below 3 pN. Again we did not observe a spatial preference of forces (Supplementary Fig. 10B).

### TCR-imposed forces exerted by cytolytic CD8^+^ OT-1 transgenic T-cells

While antigen recognition by CD4+ and CD8+ T-cells shares many underlying molecular and subcellular processes, functional consequences differ considerably. Since many studies on TCR-imposed forces have been conducted with the use of CD8+ T-cells (14, 20), we also quantitated tensile forces with the use of cytolytic T-cells (CTLs) isolated from spleen and lymph nodes of OT-1 TCR-transgenic mice and differentiated *ex vivo*. As shown in Supplementary Fig. 6B force analysis involving the use OT-1 TCR-transgenic CTLs interacting with MFS-H57 presented on gel-phase SLBs gave rise to histograms that were similar to those observed earlier with the use of CD4^+^ 5c.c7 T-cells. They yielded a high force peak of 6.1 ± 0.1 pN under activating and 5.2 ± 0.1 pN under scanning conditions.

### Temporal analysis of forces acting on single TCR-bound MFS molecules

We next determined the force kinetics of single TCR-bound MFS-H57 molecules. We exclusively analyzed MFS-H57 entities, which could be ascribed without ambiguity to the low-FRET peak, i.e. trajectories with at least five FRET efficiency data points below 0.6. The variability in the FRET signals within single traces was unexpectedly low: we observed an average standard deviation per trajectory of 0.12 (corresponding to force fluctuations of 1.5 pN), which was substantially lower than those derived from all low-FRET signals (equaling 0.22 and corresponding to 2.8 pN). This indicates that MFS-exerted forces covered a broad range but remained rather constant over the entire duration of the trajectories (on average 0.9s recorded at 50 frames per second). To investigate transitions in force amplitudes applied on individual TCRs in more detail, we conducted experiments at considerably lower frame rates with trajectories recorded for up to 71s on average. With decreasing frame rates, we observed more frequent force transitions within single trajectories. Accordingly, the dispersion of force values per single molecule trajectory increased towards the dispersion of all recorded force data. Based on this and the representative single molecule FRET trajectories shown in Fig. 3, we conclude that transitions between FRET levels occurred on a time scale of seconds.

**Figure 3:**
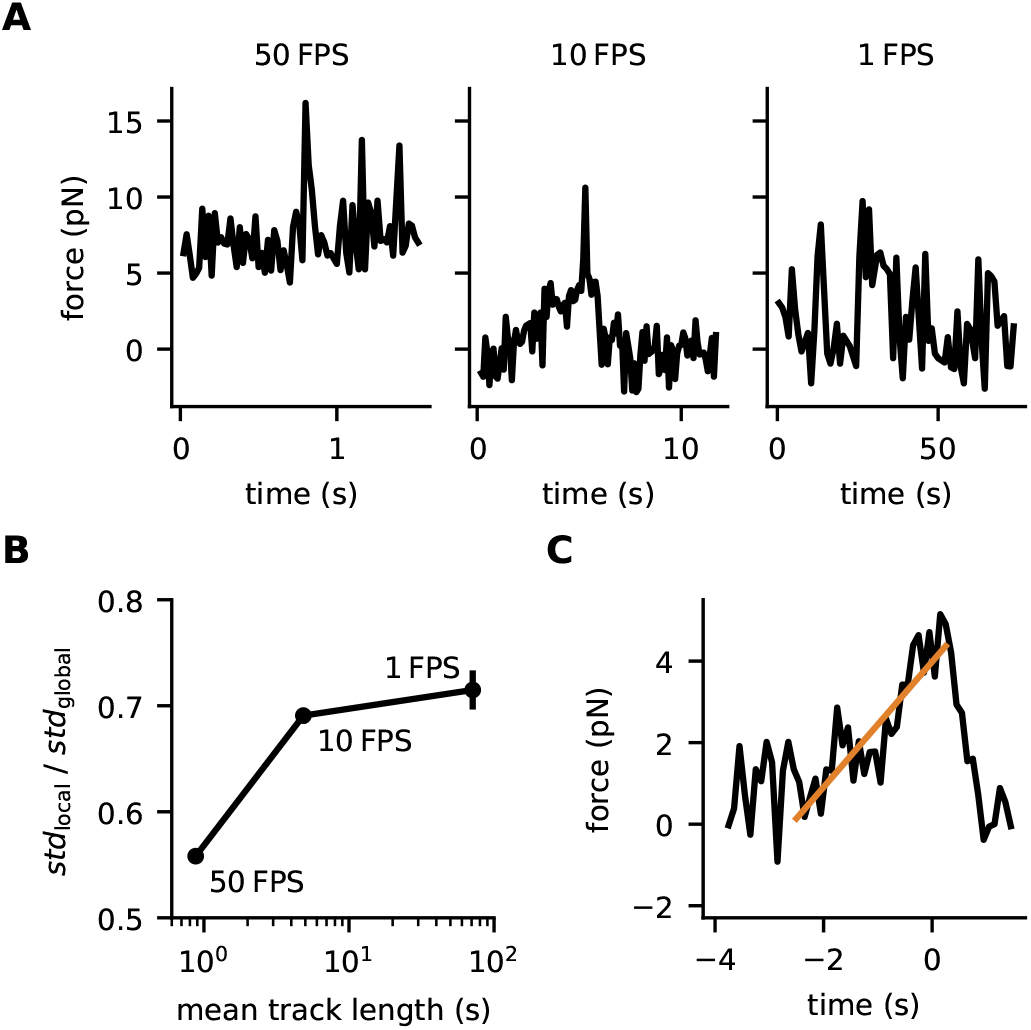
Temporal analysis of TCR-applied forces observed for activated T-cells confronted with gel-phase SLBs. (**A**) Representative single-molecule force traces recorded at indicated frame rates. We observed transitions within force trajectories only in traces recorded at low frame rates, whereas trajectories recorded at high frame rates showed rather constant force levels. (**B**) For quantitative readout, we determined the force fluctuations of each trajectory as standard deviation, calculated the average over all trajectories (termed “std_local_”), and divided the number by the global standard deviation of all measured force values (termed “std_global_”). Ergodic behavior was increasingly observed with decreasing frame rates. Error bars are standard errors of the mean. 4073 events in 94 tracks from 593 movies in dataset (50 fps), 1823 events in 38 tracks from 368 movies in dataset (10 fps), 649 events in 10 tracks from 108 movies in dataset (1 fps). (**C**) Assessment of the loading rate for single-molecular force transitions recorded at 10 fps. The panel displays a pooled analysis of 470 data points taken from 7 clearly identifiable single-molecule force ramps, which were synchronized with respect to the force peak. A linear fit (orange line) yielded an average loading rate of 1.5 pN s^-1^.

For global analysis, we pooled all clearly identifiable force ramps from several experiments by synchronizing the traces with respect to the force peak, yielding an average single molecule pulling trace (Fig. 3B). The force increased with an average loading rate of 1.5 pN s^-1^ for approximately 2.5s, before the accumulated force was released in a step-wise fashion. This behavior was reminiscent of load-fail events observed in traction-force microscopy (19), yet at 100-fold smaller peak force levels, and could be ascribed to rupture of the connection between the TCR and the cytoskeleton or between the TCR and the MFS-H57.

### TCR-imposed forces on the natural ligand pMHC

We next quantified tensile forces exerted by the TCR on agonistic pMHC. For this we employed gel-phase SLBs decorated with the second force sensor described above for which the flagelliform peptide was connected to IE^k^/MCC (MFS-IE^k^/MCC) instead of H57. Functional integrity of the construct was verified by measuring the calcium response of 5c.c7 TCR-transgenic T-cell blasts (Supplementary Fig. 3C). Activating conditions gave rise to a clearly discernible peak corresponding to mean forces of 2.0 ± 0.0 pN (Fig. 4) resulting from TCR-pMHC bonds sustaining shear and pull, at least to some degree: when compared to H57-TCR binding, the majority of IE^k^/MCC-TCR binding events showed even lower tensile forces close to the MFS-based detection limit of 0.9 pN. Such differences became even more markedly evident under scanning conditions, where we no longer observed a visible high-force shoulder. This latter finding may result from early TCR-pMHC bond rupture prior to establishing maximal force. Alternatively, it may reflect a comparatively low binding efficiency between MFS-IE^k^/MCC and the TCR under scanning conditions with few pulling events being masked under the majority of non-ligated sensors. Further studies will be required to discriminate between these possible scenarios.

**Figure 4:**
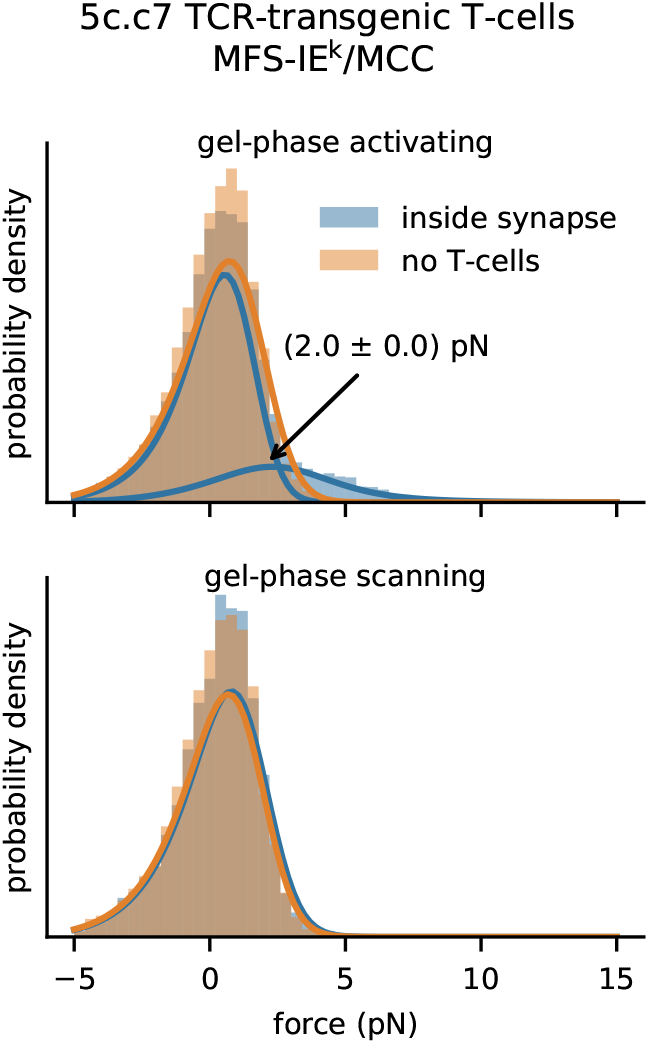
Single-molecule pulling forces exerted via single TCRs on the pMHC-functionalized molecular force sensor. Single molecule analysis of MFS-IE^k^/MCC on gel-phase SLBs recorded under activating (top) or scanning conditions (bottom), using antigen-experienced 5c.c7 TCR-transgenic T-cells. We observed a significant high force peak at 2.0 ± 0.0 pN for the activating case amounting to 24% of all events, which was no longer detectable under scanning conditions. n = 18763 (770 movies) for activated T-cells, n = 14991 (205 movies) for stimulatory SLBs without T-cells, n = 8646 (348 movies) for scanning T-cells, n = 5120 (106 movies) for non-stimulatory SLBs without T-cells.

## DISCUSSION

In recent years, forces exerted on TCR-pMHC bonds have been frequently associated with T-cell antigen recognition (4–6, 38). The origin of these forces has been linked to actomyosin-based contractions in case of migrating T-cells, and actin reorganization during synapse formation as a consequence of T-cell activation (39). Indeed, in resting cells ∼30% of the TCR are linked to the cytoskeleton (40). A clutch-like coupling between the cortical actin and microcluster-resident TCRs has been proposed for their directional transport to the synaptic center following T-cell activation (41). It is therefore conceivable that T-cells exert tensile forces onto TCR-pMHC bonds.

Our MFS-based findings are best explained by TCR-exerted forces which act largely tangentially with respect to the T-cell surface. In our T-cell stimulation regimen, fluid-phase SLBs provide an environment with substantially lower viscous drag to its embedded proteins than gel-phase SLBs. Of note, the drag *γ* experienced by a lipid-anchored protein in a fluid-phase membrane is smaller than 3×10^-9^ Pa s m (42). As a consequence, MFSs pulled with the velocity of a TCR microcluster migrating to the synaptic center (*v* = 0.14 µm s^-1^ (43)) would be subjected to a viscous force *F* that is lower than 0.4 fN (with *F* = *γ* · *v*) and therefore negligible when compared to *F_min_*. Thus, moving the MFS laterally within fluid-phase SLBs does not result in a drag that is sufficient to stretch the sensor. In contrast, the viscous drag encountered in a gel-phase membrane is large enough to arrest embedded MFSs. As a consequence, the MFS becomes invariably extended whenever the TCR places it under lateral load.

Pulling the MFS away from the SLB will result in its elongation. This is because the detectable maximum force *F_max_* is substantially lower than the force required to pull six to nine lipids associated with the mSAv-3xHis_6_ (44) out of the membrane (> 7 pN at loading rates >2 pN s^-1^ per single lipid (45) amounting to > 42 pN for six parallel anchors). Furthermore, extracting a biotin from its binding pocket requires more than 25 pN at loading rates larger than 1 pN s^-1^ (46).

Our observations are consistent with the reported occurrence of tangential forces detected via traction force microscopy (18, 19). In this study, TCR-exerted forces of 50 to 100 pN were detected per pillar with a cross-sectional area of ∼0.75µm^2^. Considering a parallel configuration of the bonds and TCR densities of approximately 70 molecules per µm^2^, as they are typical for most T-cells (47), traction forces per bond would amount to a low piconewton regime, similar to the results of our study. In contrast, Ma and colleagues reported TCR-imposed forces of 4.7 pN when T-cells contacted fluid-phase SLBs (21). However, in that study multiple DNA tension probes, and, as a consequence, multiple TCR molecules were cross-linked via the solid gold nanoparticles, allowing for tensile forces between the cross-linked TCRs.

Of particular importance, we observed no indication that molecular forces beyond 2 pN exerted via the TCR would be a consequence of or a precondition for T-cell signaling:

(i) We detected a high-force peak exclusively when T-cells had been seeded onto gel-but not fluid-phase SLBs, regardless of whether T-cells had been stimulated in the course of contact formation or not. For scanning T-cells, we attribute such forces to the contractility of the actomyosin cytoskeleton, to which plasma membrane-resident TCRs are coupled.

(ii) Furthermore, our results render it unlikely that forces larger than 2 pN and imposed on the TCR are required to trigger the TCR. After all, T-cells seeded onto functionalized fluid-phase SLBs did not give rise to changes in FRET of interacting MFS corresponding to forces higher than 2 pN even though they showed all the hallmarks of TCR-proximal signaling, such as elevated intracellular calcium levels, recruitment of ZAP70 kinase, TCR-microcluster formation, and TCR-movement towards the synaptic center. It should be noted that at this stage we cannot rule out the presence of a minority of molecules which may have experienced higher tensile forces at the onset of T-cell triggering and which would have been missed by our analysis. However, even in this hypothetical case contributions from tangential forces can be ruled out. While MFS-H57 was specifically designed in our study for its capacity to maintain TCR-bonds even under TCR-mediated pulling, nominal pMHCs are typically characterized by fast TCR-dissociation requiring a significantly lower activation barrier for bond rupture: In case of IE^k^/MCC binding to its cognate 5c.c7 TCR, already forces in the lower pN-regime appear likely to induce the dissociation of pMHC from the TCR binding pocket, thereby reducing the average bond lifetime (4). If TCR-pMHC dissociation is promoted by acting forces, force loads may no longer materialize to the same magnitude, as they are exerted in case of stable TCR-antibody interactions. Our observation of reduced pulling forces in case of MFS-IE^k^/MCC would hence be in agreement with the presence of slip bonds formed between the 5c.c7 TCR and IE^k^/MCC, as was previously reported (9).

Interestingly, our time-resolved experiments did not reveal constant but linearly increasing forces (Fig. 3C). Such force ramps not only affect the mean but particularly the distribution of bond lifetimes, an effect which could be exploited by T-cells to improve the specificity and sensitivity of antigen discrimination (4, 48). Previously, we have reasoned by analytical deduction that loading rates of only 1 pN/s would already suffice to enable peptide discrimination with high specificity and sensitivity (4). Taken together, our observations comply with a slip bond behavior for the interaction between the 5c.c7 TCR and IE^k^/MCC. However, we cannot completely rule out catch bond formation at this stage, if such bonds feature a transition to slip bonds at forces below ∼1 pN.

T-cells scan the surface of the opposing antigen-presenting cell (APC) typically via microvilli-like protrusions (27), a process that has also been described for T-cells establishing a contact with activating surfaces (27, 49, 50). During this phase, the limited number of TCRs that can bind pMHC on the opposing APC become subjected to tangential and perpendicular forces upon pMHC binding and until TCR-ligand bond rupture. For tangential forces to act on pMHC-engaged TCRs within the cellular interface, MHC molecules need to be subjected to a counterforce at the level of the APC membrane, as could be inferred from the rather considerable number of immobile or slowly diffusing MHC II molecules residing in the plasma membrane of dendritic cells of ∼50% (51). Since the perpendicular forces measured here on fluid membranes via MFS-H57 are similar in magnitude to the average forces experienced by the MFS-IE^k^/MCC, we consider it likely that even rather weak pulling forces exerted in perpendicular direction would suffice to terminate TCR-pMHC bonds. If T-cells fail to become activated while probing the APC surface, T-cells reduce pulling forces as is indicated by our temporal analysis (Supplementary Fig. 9). Reduced pulling forces over time may reflect an adaptation towards a, e.g. more sensitive scanning process, possibly at the costs of diminished selectivity. Antigen-dependent and -independent tonic TCR-mediated signaling, which has been reported to instruct T-cell-fate decisions and provide survival signals, may well depend on lowering the force regime.

A variety of mechanistic and phenomenological models (52, 53) have been proposed to explain how extracellular TCR-pMHC binding is translated with high sensitivity into a ligand-specific intracellular signaling event. With rising intracellular calcium levels increase, LFA-1 undergoes an affinity switch to consolidate via ICAM-1-binding T-cell adhesion with the rapid formation of TCR-microclusters in the synaptic periphery. Within a microcluster multiple TCR-pMHC bonds are placed under load in parallel. Our study points at consequences that differ for perpendicular and tangential pulling directions. Under activating conditions forces were not detectable in perpendicular direction, possibly reflecting force dissipation over the parallel bonds. When T-cells were confronted with gel-phase SLBs, forces imposed on individual TCRs appeared however similar under both activating and non-activating conditions, implying a different mechanism of force exertion in tangential direction. Microclusters are concomitantly transported tangentially towards the synaptic center with a broad range of velocities amounting to 20 ± 8 nm s^-1^, a process that was reported to be strongly affected by constraints built into the SLB (41, 54): TCR-microclusters encountering barriers in an orientation perpendicular to their movement were found arrested, while barriers positioned in intermediate angles decelerated transport (41). The latter finding can be rationalized by the loose clutch mechanism of TCR interacting with the cytoskeleton (41), thereby spreading the average load over time in a make-and-break cycle. We consider it plausible that our single molecule force traces shown in Fig. 3 represent such transient links and that the properties of our gel-phase SLBs prevented the transport of ligated TCR along the T-cell surface in analogy to the previously described diffusion barriers. Indeed, after force had accumulated for ∼ 2.5 s, the connection between the TCR and cytoskeletal drag appeared to be released, resulting in an immediate drop in the force applied to the MFS. The sensor’s stretching rate of 4 nm s^-1^ as calculated from the observed loading rate of 1.5 pN s^-1^ (for more details see Methods section, force sensor calibration, Eq. 3) corresponds to the lower end of the reported spectrum for microcluster transport rates, which probably reflects reduced cytoskeletal dynamics under load. The delayed rise in pulling forces observed for T-cells encountering activating gel-phase membranes (Supplementary Fig. 9A) further supports this interpretation, as microclusters need to be formed beforehand. Further experiments will be needed to address the mechanical origin of the linear load increase. In agreement with previous findings (41) our data rules out a direct coupling to retrograde actin flow, since the observed stretching rates of our sensors of a few nanometer per second do not comply with actin flow rates of several tens of nanometers per second (55).

In summary, we found that TCR-mediated forces occur within the immunological synapse in the 4-9 pN regime only when TCR-ligands were embedded in a gel-phase SLB, indicating their predominantly shear force nature. Forces exerted on the natural ligand pMHC were of substantially smaller magnitude, and may indicate force-accelerated rupture of the TCR-pMHC interaction. Given the high spatiotemporal resolution afforded, we expect the experimental system presented in this study to prove highly adequate and versatile to further studies delineating the role and characteristics of TCR-imposed forces in TCR-proximal signaling and ligand discrimination.

## MATERIALS AND METHODS

### Force sensor - anchor unit

Recombinant mSAv-3xHis_6_ served as anchor unit for the force sensor and was produced as described (30). Briefly, biotin binding deficient subunits equipped with a C-terminal His_6_-tag – “dead” – and functional subunits with C-terminal 3C protease cleaving site followed by Glu_6_-tag – “alive” – were expressed as inclusion bodies in E.coli BL-21 using the pET expression system (Novagen). Inclusion bodies were dissolved in 6 M Guanidine hydrochloride at 10 mg/ml, mixed at a ratio of 3 “dead” to 1 “alive” subunit and refolded in 100 volumes PBS. The salt concentration was reduced to 2 mM by cycles of concentration using a 10 kDa cutoff spin concentrator (Amicon) and dilution with 20 mM TRIS pH 7.0. The mixture was loaded onto a MonoQ 5/50 column (GE Healthcare), and after washing the column with 100mM NaCl in 20 mM TRIS pH 7.0 mSAv-3xHis_6_ was eluted at 20 mM TRIS pH 7.0, 240 mM NaCl. Human 3C protease (Pierce) was added and allowed to cleave the Glu_6_-tag overnight. The digested protein was subjected to Superdex 200 (GE Healthcare) size exclusion chromatography in PBS to yield to pure anchor unit.

### Force sensor - spring unit

The MFS peptide biotin-N-GG<u>C</u>GS(GPGGA)_5_GG<u>K</u>YGGS-K(ε-N_3_) (residues for fluorophore attachment are underlined), MFS_C_ peptide biotin-N-GGGGS(GPGGA)_5_GG<u>K</u>Y<u>C</u>GS-K(ε-N_3_) and the MFS_0_ peptide biotin-N-GGGGS (GPGGA)_5_ GGKYG GS-K(ε-N_3_) were purified via preparative C18 reverse phase HPLC (5 µm particle size, 250×21.2 mm, Agilent) applying a linear gradient of 0.1% triflouracetic acid (TFA) in deionized water (buffer A) and 0.1 % TFA, 9.9 % deionized water, 90 % acetonitrile (buffer B) over 80 minutes at a flow rate of 5 ml/min. The identity of the spring modules was confirmed via MALDI-TOF mass spectrometry (Bruker). 350 µg lyophilized peptide was dissolved in PBS at a final concentration of 5 mg/ml, mixed with Alexa Fluor 555 maleimide (Thermo Fisher Scientific) at a 1.5 fold molar surplus and then incubated for 2 hours followed by an incubation with 1.5-fold excess Alexa Fluor 647 succinimidyl ester (Thermo Fisher Scientific) in the presence of 330 mM NaHCO_3_ for 2 hours at room temperature. HPLC purification and lyophilization was performed after each step and successful conjugation was confirmed via mass spectrometry.

### Force sensor - functional unit

TCRβ-reactive H57 antibody single chain fragments featuring a C-terminal unpaired cysteine residue (H57 cys scF_V_) were expressed in E.coli BL-21 as inclusion bodies using the pET expression system (Novagen) and refolded as described (9, 40). The product was then concentrated using a 10 kDa cutoff spin concentrator (Amicon) in the presence of 25 µM tris-(2-carboxyethyl)phosphine hydrochloride (TCEP, Pierce). Monomeric H57 cys scF_V_ was purified by Superdex 75 (GE Healthcare) size exclusion chromatography in PBS, then concentrated to 1 mg/ml in the presence of 25 µM TCEP and allowed to react with a 10-fold surplus of dibenzylcyclooctyne-maleimide (DBCO, Jena Bioscience) for 2 hours at room temperature. Unreacted DBCO was removed by gel filtration and proteins were brought to a concentration of 1 mg/ml.

IE^k^ α featuring an additional C-terminal cysteine residue and IE^k^ β were also expressed as inclusion bodies and refolded in the presence of MCC 88-103 peptide (ANERADLIAYLKQATK, Intavis) for 2 weeks as previously described (9). Correctly folded IE^k^ molecules were isolated via an in house built 14-4-4 antibody column, reduced with 25 µM TCEP followed by S200 gel filtration. Monomeric IE^k^ MCC with unpaired cysteine residue was collected and coupled to DBCO as described above.

### Force sensor assembly

Refolded from E.coli inclusion bodies H57-scF_V_ were modified for sensor attachment with DBCO maleimide at a mutant unpaired C-terminal cysteine residue (Supplementary Fig. 1A-D). MFS peptides were mixed with DBCO conjugated proteins and 25 Vol % 2 M TRIS pH 8.0 at a molar ratio of at least 5:1 and incubated for 2 hours at room temperature. Unreacted protein was removed with the use of a monomeric avidin column (Pierce). After elution with biotin, unconjugated spring was separated from MFS conjugates using size exclusion chromatography on a Superdex 75 column (GE Healthcare). mSAv-3xHis_6_ gel shift assays documented successful conjugation as only a neglectable proportion of H57-scFv gave rise to dimers (Supplementary Fig. 1E). The existence of such entities may have yielded a small underestimation in ensemble readouts. However, in the case of single-molecule experiments, multimeric structures could be directly identified and subsequently eliminated from the analysis.

### T-cells

Antigen-experienced 5c.c7 murine T-cell blasts were obtained as previously described (56). Briefly, 7.5*10^6^/ml spleenocytes were stimulated with 2 µM HPLC-purified MCC peptide (ANERADLIAYLKQATK, Intavis Bioanalytical Instruments) in T-cell medium, which consisted of RPMI 1640 (Gibco) supplemented with 10 % FCS (Sigma), 100 U/ml penicillin/streptomycin (Gibco), 2 mM glutamate (Gibco), 1mM sodium pyruvate (Gibco), 1x non-essential amino acids (Gibco) and 50 µM ß-Mercaptoethanol (Gibco). Culture volume was doubled and also supplemented with 100 U/ml IL-2 (eBioscience) on day 2. T-cell cultures were expanded in a ratio of 1:1 on day 3 and 5. Dead cells were removed by centrifugation on a cushion of Histopaque-1119 (Sigma) on day 6. T-cells were used for experiments on day 7-9 after initial stimulation.

OT-1 aβ TCR-transgenic mice of 8-12 weeks were sacrificed to collect spleen and lymph nodes. Isolated splenic antigen-presenting cells were loaded with 1 µM C18 reverse-phase HPLC–purified Ovalbumin (Ova 257–264, SIINFEKL) peptide. After 1 hour incubation, splenic antigen-presenting cells were washed with RPMI 1640 culture medium (Gibco). Subsequently, lymph-node derived cells were mixed in a 1:2 ratio with OVA-peptide pulsed spleenocytes and cultured in RPMI 1640 culture medium (Gibco) supplemented with 10% fetal calf serum (FCS) (Sigma), 100 U/ml penicillin/streptomycin (Gibco), 2 mM glutamate (Gibco), 1 mM sodium pyruvate (Gibco), 1x non-essential amino acids (Gibco) and 50 µM ß-mercaptoethanol (Gibco) at 37°C and 5% CO2. On day 2, IL-2 was added to a final concentration of 50 U/ml. Cells were split 1 :1 with fresh media every second day (day 3, 5 and 7). Dead cells were removed on day 7 prior to splitting by Ficoll-Paque Plus (Merck) step gradient centrifugation and cytotoxic T lymphocytes (CTLs) were used on day 8-10 for experiments.

### Ethical compliance statement

Animal were bred and euthanized for T-cell isolation according to guidelines and protocols evaluated by the ethics committee of the Medical University of Vienna and approved by the Federal Ministry of Science, Research and Economy, BMWFW (BMWFW-66.009/0378-WF/V/3b/2016). Animal husbandry and experimentation were performed under national laws (Federal Ministry of Science, Research and Economy, Vienna, Austria) and the ethics committee of the Medical University of Vienna and according to the guidelines of the Federation of Laboratory Animal Science Associations (FELASA).

### Reagents

Hank’s buffered salt solution (HBSS) and phosphate buffered saline (PBS), Cytochalasin D (from Zygosporium masonii) as well as fetal calf serum (FCS) were from Sigma Aldrich (Merck KGaA, Germany). DPPC (1,2-dipalmitoyl-sn-glycero-3-phosphocholine), POPC (1-palmitoyl-2-oleoyl-glycero-3-phosphocholine), and Ni-NTA-DGS (1,2-dioleoyl-sn-glycero-3-[(N-(5-amino-1-carboxypentyl)iminodiacetic acid)succinyl] (nickel salt)) were from Avanti Polar Lipids, Inc., USA. ICAM-1 and B7-1 were produced in Hi-5 insect cells using baculovirus and purified as described via size exclusion and ion exchange chromatography (9).

### Preparation of Functionalized Lipid Bilayers

98 mol-% POPC or DPPC and 2 mol-% Ni-NTA-DGS, all dissolved in chloroform, were mixed and then dried under nitrogen flow for 20 min. The lipid mixture was suspended in 1 ml PBS at room temperature (POPC) or 55 °C (DPPC) and then sonicated for 10 minutes in an ultrasound bath (USC500TH, VWR, England) at the same temperature to form small unilamellar vesicles (SUVs). The SUV solution was finally diluted to 125 μM using PBS.

Cover slips (MENZEL-Gläser Deckgläser 24 × 60 mm #1.5) were treated in a plasma cleaner (PDC-002 Plasma Cleaner Harrick Plasma, Ithaca, NY, USA) for 10 minutes and glued to the bottom of an 8-well chamber (Nunc Lab-Tek, Thermo Scientific, USA) using duplicating silicone (Twinsil soft 18, picodent, Germany). SUVs were incubated for 20 minutes at room temperature (POPC) or 55 °C (DPPC) to form bilayers. After incubation, excess vesicles were washed away using PBS.

His-tagged proteins (for activating conditions: 10 ng mSAv-3xHis_6_, 30 ng ICAM-1, 50 ng B7-1; for scanning conditions: 0.4 ng mSAv-3xHis_6_, 30 ng ICAM-1) were added in 0.5 ml PBS to each well and incubated for 75 minutes at room temperature. Afterwards, the bilayer was washed with PBS.

Finally, the bilayer was incubated for 20 minutes with biotinylated MFS variants (for activating conditions: 10 ng MFS_0_, 5 – 20 pg MFS or MFS_C_; for scanning conditions: 20 – 60 pg MFS or MFS_C_) in 0.5 ml PBS per well for binding to the mSAv-3xHis_6_. Bilayers were washed again to remove excess MFS. Immediately before adding cells, the buffer was exchanged for HBSS containing 2% FCS.

For ensemble experiments, molecular densities were determined by dividing the fluorescence signal per pixel by the single molecule brightness recorded at the same settings, considering the effective pixel width of 160 nm. In single molecule samples, densities were determined by counting the number of molecules and dividing by the area of the field of view.

### T-cell calcium imaging

3 ml of HBSS + 2% FCS were added to ∼10^6^ T-cells. After centrifugation (300 RCF for 3 min), supernatant was removed and cells were incubated with 1 μg of Fura-2 AM dye (Thermo Scientific, USA) at 1mg/ml DMSO in a total volume of 100µl for 20 minutes at room temperature, and washed (addition of 5 ml HBSS + 2% FCS, centrifugation at 300 RCF for 3 min, removal of supernatant, resuspension in 100µl). Cells were kept at room temperature until seeding them onto the functionalized bilayers and for a maximal duration of 30 minutes.

Time-lapse movies were recorded using alternating excitation (340 and 380 nm, 50 and 10 ms illumination, respectively) using a monochromatic light source (Polychrome V, TILL Photonics), coupled into a Zeiss Axiovert 200M microscope equipped with a 10x objective (UPlanFL N 10x/0.30, Olympus) and an Andor iXon Ultra 897 EMCCD (Andor Technology Ltd, Belfast, UK) camera. The total recording time was 15 minutes or longer at 1 Hz.

Cells were tracked in each frame using a particle tracking algorithm published by Gao and Kilfoil (57). Tracking parameters were chosen accordingly to only include cells in contact with the SLBs. A custom-built MATLAB software was used to generate the ratio images for each frame. We used the average ratio of each cell in a central circle of 2µm.

Population analysis was performed using a custom-build MATLAB software based on the “Methods for Automated and Accurate Analysis of Cell Signals” (58). Briefly, all ratio traces of a population were normalized frame-wise to the population median of the negative control. Cells with at least 80% of their trajectory above activation threshold were counted as activated. Individual trajectories were aligned to the maximum ratio value of each trajectory creating a synchronized population with respect to peak position. The population was synchronized with respect to cell adhesion and contained individual cellular trajectories starting from cell contact with the SLB. The data was displayed as medians with interquartile range. As positive control we used SLBs decorated with 30 ng ICAM-1, 50 ng B7-1 and 10 ng His-tagged MCC-IE^k^ amounting to ∼100 molecules µm^-2^, as negative control SLBs decorated with only ICAM-1.

### T-cell fixation and immunostaining

Prior to T-cell seeding, ligand densities on SLBs were determined as described above. 5c.c7 TCR-transgenic T-cells were seeded onto SLBs and allowed to settle for 10 minutes at room temperature. Fixation buffer (PBS, 8% formaldehyde (28908, ThermoFisher Scientific), 0.2% glutaraldehyde (G7776-10ML, Sigma Aldrich), 100 mM Na_3_O_4_V (S6508-10G, Sigma Aldrich), 1 M NaF (S7920-100G, Sigma Aldrich)) was added 1:1 and incubated for 20 minutes at room temperature, before samples were rinsed with PBS. Permeabilization buffer (PBS, 100 mM Na_3_O_4_V, 1 M NaF, 0.2% Triton-X (85111, ThermoFisher Scientific)) was added and after 1 minute samples were washed with PBS before blocking with passivation buffer (PBS, 3% BSA) for 30 min. Anti-ZAP70 antibody labeled with AF647 (clone 1E7.2, #51-6695-82, ThermoFisher Scientific) was added at a final concentration of 1.25 μg ml^-1^ and incubated overnight at 4°C. Samples were washed extensively with passivation buffer. Imaging was performed in HBSS.

### Inhibition of actin polymerization

5c.c7 T-cells were seeded onto SLBs in HBSS containing 2% FCS and incubated for 7 minutes at room temperature. 10 μM Cytochalasin D in DMSO was carefully added and was incubated for 10 minutes at room temperature prior to single-molecule FRET measurements. A mock sample was incubated with the same volume of DMSO. Effect of inhibition was monitored in parallel by calcium flux measurements.

### Microscopy Experiments

Experiments were performed at room temperature and carried out on two different microscopy systems. *Microscopy system #1:* Fluorophores were excited with 640nm (OBIS 640, Coherent, CA) or 532 nm (Millenia Prime, Spectra Physics, CA) laser light coupled into an epifluorescence microscopy (Zeiss, Germany). All experiments were performed in objective-type total internal reflection (TIR) configuration, using a high-NA objective (α Plan-FLUAR 100x/1.45 oil, Zeiss, Germany) and a quad-band dichroic mirror (Di01-R405/488/532/635-25×36, Semrock Inc., USA) to separate excitation light from emission light. The fluorescence emission path was split along wavelengths with the use of an emission beam splitter system (Optosplit II, Cairn Research, Kent, UK) employing a dichroic mirror with a cutoff wavelength of 640nm (FF640-FDi01-25×36, Semrock Inc., USA) and emission filters (ET570/60m and ET675/50m band pass, Chroma Technology Corp, USA). Both color channels were imaged side-by-side on the same chip of an EM-CCD camera (Andor iXon Ultra 897, Andor Technology Ltd, UK). In addition, we used a 405 nm laser (iBeam Smart 405-S, Toptica Photonics AG, Germany) for visualizing the cell contours.

*Microscopy system #2:* Excitation was achieved by coupling a 532nm (OBIS LS, Coherent, USA) and a 640 nm (iBeam smart, Toptica Photonics, Germany) laser line into a T*i*-E inverted microscope (Nikon, Japan) via a dichroic mirror ZT405/488/532/640rpc (Chroma, USA) into a 100x objective (SR Apo TIRF, Nikon, Japan) in an objective-based TIR setting. Emission was split using an Optosplit II (Cairn Research, UK) equipped with a 640 nm dichroic mirror (ZT640rdc, Chroma, USA) and emission filters (ET575/50, ET655LP, Chroma, USA) and simultaneously imaged on an Andor iXon Ultra 897 EM-CCD camera (Andor Technology, UK). Calcium imaging was performed using a Lambda LS xenon lamp (Sutter Instruments, USA) with excitation filters (340/26, 387/11, Semrock, USA) for illumination via a 20x objective (S FLUOR, Nikon, Tokyo, Japan) and Fura-2 emission filter (ET525/50, Chroma, USA). Devices were controlled by MetaMorph imaging software (Molecular Devices, USA).

Microscopy system #1 was used for all single molecule experiments and for ensemble FRET measurements with the use of DPPC bilayers. Microscopy system #2 was employed for ensemble FRET experiments using POPC membranes and for calcium measurements.

### Ensemble FRET measurements

FRET efficiencies were determined at the ensemble level using donor recovery after acceptor photobleaching (DRAAP). For this, an image of the donor channel was recorded before and after acceptor photobleaching, and the pixel-averaged fluorescence signal *f*pre and *f*post, respectively, was calculated for each cell. The FRET efficiency was then given by *E*bulk = (*f*post -*f*pre)/*f*post. For quantitation an area was selected underneath the T-cell and averaged, giving rise to 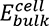. The same area was also recorded on lipid bilayers devoid of T-cells, analyzed for FRET efficiency, and averaged, yielding 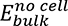. Finally, the difference was calculated via 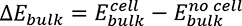. Bleaching times were between 500ms and 1s, illumination times and delays were typically below 10ms. Single-molecule FRET measurements

### Single-molecule FRET measurements

In a typical experiment we recorded 300 images alternating between green and red excitation, with illumination times t_ill_=5ms (see Supplementary Fig. 5B). The delay between the green and the red color channel, t_gr_, was kept as short as possible, typically t_gr_=5-15ms. The delay between two consecutive green-red doublets, t_delay_, was chosen in a range between 5ms and 975ms. For each time point, three images were recorded: donor emission upon donor excitation (*Im*DD), acceptor emission upon donor excitation (*Im*DA), and acceptor emission upon acceptor excitation (*Im*AA). To convert camera counts into emitted photons, count values were multiplied by the manufacturer-provided camera efficiency at the chosen settings (15.7 photons per count) and divided by the EM gain (typically, 300).

To determine single-molecule FRET efficiencies, a series of seven processing steps was performed (Supplementary Fig. 5 and Supplementary Fig. 11). Data analysis was implemented in Python 3, using the Numpy, SciPy (59) and Pandas packages for general numerical computations, matplotlib for plotting and path handling (60), scikit-learn for fitting of Gaussian mixture models (61), trackpy for single-molecule tracking (62), pims for reading image sequences from disk, and OpenCV for adaptive thresholding. Jupyter notebook and ipywidgets were used as a front-end to Python.

#### 1. Image registration

Two-color images (*Im*DD and *Im*DA) were registered and corrected for chromatic aberrations using beads of 200nm diameter (TetraSpec, Invitrogen) as fiducial markers, by fitting an affine transformation to the set of localization pairs. In our case, we transformed and interpolated the donor color channel to match the acceptor color channel (*T*(*Im*DD)).

#### 2. Identification of candidates for single-molecule signals

We localized single-molecule signals on the sum image *T*(*Im*DD) + *Im*DA and on the image *Im*AA (63), and interlaced the resulting localizations according to the recording sequence. Taking the sum image ensured the detection of both low FRET and high FRET signals with high sensitivity. The resulting single-molecule position data were linked into trajectories using a single-molecule tracking algorithm (62), with a maximum search radius of 800nm between two consecutive frames. Only trajectories with a length 2: 4 observations were considered. We allowed for gaps of up to three frames in the trajectories to account for bleaching and blinking of fluorophores as well as missed signals. These gaps were filled using linear interpolation between the corresponding positions before and after the missed events. As a result, we obtained the coordinates and according trajectories of all candidates for single-molecule signals in the coordinate system of the acceptor channel. For further processing, the coordinates corresponding to signals recorded in the donor channel were transformed back to the coordinate system of the donor channel via *T*^-1^.

#### 3. Determination of fluorescence brightness values

We first applied a Gaussian blur (σ = 1 pixel) to the three raw data images *Im*DD, *Im*DA, and *Im*AA, which reduces noise while preserving the signal intensity (Supplementary Fig. 5D). To determine the single-molecule brightness, the camera counts in a 9px-diameter foreground circle around each signal’s position, s_ij_, were considered (Supplementary Fig. 5C). Local background, bg, was estimated by calculating the mean camera count in a ring around the signal circle (excluding any pixels affected by nearby signals). Fluorescence brightness was calculated according to *f* = ∑i,j∈circle(*s*ij -*bg*). Finally, f was corrected for uneven excitation laser profile across the field of view *f* ⟼ *f*(*x*, *y*)⁄*profile*(*x*, *y*). We determined *profile*(*x*, *y*) for each laser from heavily stained samples, applied a Gaussian blur (σ=10px), and scaled to a maximum value of 1. For each signal, we hence obtained here three values: the fluorescence brightness at donor excitation in donor emission (*f*DD), donor excitation in acceptor emission (*f*DA), and acceptor excitation in acceptor emission (*f*AA).

#### 4. Initial filtering steps

We applied several filter steps to disregard signals which do not match stringent criteria for single-molecules. i) remove all signals from areas with <50% of the maximum laser intensity in the donor channel. ii) remove all trajectories that show potential overlap with other trajectories. Trajectories, where >25% of data points showed overlapping foreground circles, were completely disregarded. In all other trajectories, only single signals with overlapping foreground circles were removed. iii) remove all trajectories which are not present in the first frame. This step ensures that the full photobleaching profile of each trajectory is available in the subsequent steps. iv) single step transitions were identified independently in the time trace of f_AA_ and of f_DD_+f_DA_ (64). Only trajectories were selected for further processing where f_AA_ showed single step photobleaching. Also, trajectories showing partial photobleaching in f_DD_+f_DA_ were discarded.

#### 5. Determination of stoichiometry factor and FRET efficiency

We calculated the corrected single-molecule FRET efficiency *E* and the FRET stoichiometry factor *S* according to Ref. (37) and (65):

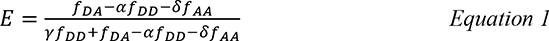

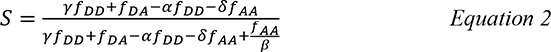

The bleedthrough coefficient, α, and the coefficient describing direct acceptor excitation, δ, were determined from separate experiments using donor only and acceptor only. The detection efficiency, γ, was determined globally from single step photobleaching events according to Ref. (66) using MFS_C_ datasets without cells as they provide large intensity differences due to high FRET efficiencies. The excitation efficiency factor β scales f_AA_; it is adjusted to reach S=0.5 in case of a calibration sample of known stoichiometry (67). In our case, we used MFS without cells as calibrations standard.

#### 6. Final filtering steps

After the correction, we applied further filtering steps. i) only data were taken before the bleaching step of the acceptor (or donor, if donor bleaching happened prior to acceptor bleaching). ii) to exclude incorrectly synthesized force sensors or clusters in the data analysis, we removed trajectories where more than 25% of the corresponding data showed S>0.6 (excluding two or more donors per acceptor) or S<0.35 (excluding two or more acceptors per donor). iii) for experiments with cells, we selected only data points underneath an adhered T-cell. We identified T-cells from Fura-2 images, which were recorded synchronously to the FRET signals, using an adaptive thresholding algorithm.

#### 7. Fitting of the FRET histograms

2D S-E data were fitted with a maximum likelihood fit, assuming a Bayesian Gaussian mixture model (61). In case of a low FRET peak exceeding the maximum detectable FRET value E_max_ (see subsection “Dynamic range of the force sensor”) or if its relative weight was smaller than 0.1 the results of a single component Gaussian fit were shown.

#### 8. Determination of fitting errors

For determination of the standard error we used bootstrapping analysis. Briefly, 100 samples (identical in size to the original data) were drawn with replacement and fitted. Specified error bars correspond to the standard deviations of the fit results.

### Calibration of the force sensor

The calibration of the force sensor is based on the study by (26), who found for the same flagelliform protein a linear force-distance relationship according to

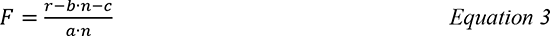

where r is the separation of the two fluorophores, b=0.044 nm the average length of a single amino acid in the collapsed state, c a constant linker length, and a=0.0122 nm pN^-1^ the compliance per amino acid. n=25 was the number of amino acids between the two fluorophores (in our case n=29). From the r-dependence of the FRET efficiency,

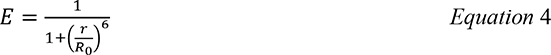

we determined the functional dependence of F on E. Here, R_0_=5.1nm denotes the Förster distance for the FRET pair AlexaF647 and AlexaF555 (68). The parameter c in Eq. 3 can be determined from experiments with F=0, which were realized here by recording MFS on a supported lipid bilayer without cells; from the according FRET efficiency E, we calculated c=2.4 nm.

Throughout the paper, we specify the expectation value of the fit results for the single molecule force distributions, which corresponds to the mean value of the observed pulling forces.

### Determination of the loading rate

Single-molecule trajectories recorded on DPPC bilayers under activating conditions at 10fps were selected for transitions between low and high force levels, based on visual inspection (see Fig. 3A, second panel for a representative example). The selected trajectories were aligned along the time axis such that the peak forces coincided, and the rising phase was fitted with a linear function. For display, single-molecule forces were averaged for each time point.

### Dynamic range of the force sensor

Simulations were implemented in Python 3. The NumPy package was used for general numerical operations as well as for random number generation.

To assess the dynamic range of the force sensor, we based our calculations on the reported elastic properties of the flagelliform protein (26). As a conservative estimate of E_min_, we considered the MFS to follow a linear force-extension relationship up to 10 pN tensile forces. Using Eq.3 and Eq.4 this relates to a minimum FRET efficiency E_min_=0.11.

E_max_ is limited by the difficulty to discriminate the low-FRET peak from the high-FRET peak. To quantify E_max_, we simulated Gaussian-distributed random data sets for the high-FRET peak, taking a mean FRET efficiency of 0.87 and a standard deviation σ=0.12, as typically observed throughout the experiments. For the low FRET peak, we simulated varying values for the mean FRET efficiency and the standard deviation at a weight of 0.15 or 0.25; the weight of 0.25 reflect the experimental data obtained with MFS-H57 on fluid-phase membranes under scanning conditions (Supplementary Fig. 7), the weight of 0.15 is a conservative estimate for a challenging scenario. Samples consisting of 2300 data points were simulated and compared to a distribution lacking the low FRET peak, using a two-sample Kolmogorov-Smirnov test.

The determined p-values are shown in Supplementary Fig. 5A (corresponding values in Supplementary Table 1): regions of low p-values correspond to parameter settings which allow for identification of a low FRET peak. As expected, the smaller the weight of the low FRET becomes, the more difficult is its identification. As a conservative estimate we took for the estimate of the dynamic range a weight of 0.15, yielding a threshold of approximately E_max_=0.8, which corresponds to F_min_=0.9 pN.

### Simulation of Single-molecule Signals

To assess the quality of the Gauss blur (Supplementary Fig. 5D), we simulated Gauss-distributed signals on a grid, with the σ-width of the Gauss being identical to the pixel size. On each pixel, the obtained number was Poisson-distributed (to simulate shot noise), and camera noise was added by a normal distributed value, which was taken from recordings on our microscopy setup. The obtained signals were analyzed as described above, both with and without the Gauss blur.

### Diffusion Analysis

To obtain diffusion coefficients in Supplementary Fig. 2A, image sequences were acquired at an illumination time of t_ill_ = 5 ms and a delay of t_delay_ = 15 ms, 95 ms or 995 ms. Single-molecule positions were determined as for FRET analysis. Mean square displacement analysis was performed as described in (69) and fitted with the equation *MSD* = 4*Dt*lag + *offset*, where D specifies the lateral diffusion coefficient, t_lag_ the analyzed time-lag, and *offset* the localization precision.

### Analysis of the mobile fraction

To determine the immobilized molecules (Fig. 2B and Supplementary Fig. 8), we calculated the smallest enclosing circle for each trajectory using the Python implementation from Project Nayuki (https://www.nayuki.io/page/smallest-enclosing-circle). Since diffusion coefficients scale with the radius of this circle, r_c_, over the square-root of the length of the trajectory, 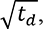 we plotted histograms of 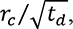 yielding peaks for the immobilized molecules 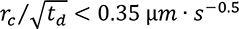 and mobile molecules 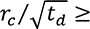 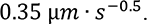

### Statistical analysis

Statistical analysis was performed with Mann Whitney U tests to evaluate differences in the mean values of the distributions. P-values <0.01 were considered as significant, ≥0.01 as not significant (n.s.). In the figure legends, we provide the number of the analyzed movies contributing to the corresponding data set, where each movie comprises between 1 and 4 T-cells.

### Code availability

We provide the Python code for single-molecule FRET analysis (https://github.com/schuetzgroup/fret-analysis) as well as the underlying Python library (https://github.com/schuetzgroup/sdt-python).

## Acknowledgements

This work was supported by the Austrian Science Fund (FWF) projects P30214-N36 (LS, GJS) and P 25775-B2 (FK, JBH), the Boehringer Ingelheim Fonds (RP), Long-Term Fellowship from Federation of European Biochemical Societies (EK), Human Frontiers Science Program RGY0065/2017 (EK), and by the Vienna Science and Technology Fund (WWTF) LS13-030 (JG, FK, LS, JBH, GJS).

## Author Contributions

JG and FK synthesized constructs. JG, FK and LS performed imaging experiments. LS developed single-molecule data analysis tools. HS and RP provided important insights. G.J.S, J.B.H. and E.K. conceived the study. JG, FK, LS, JBH and GJS contributed to data analysis, interpretation of the results and wrote the manuscript.

## SUPPLEMENTARY FIGURE LEGENDS

**Supplementary Figure 1:**
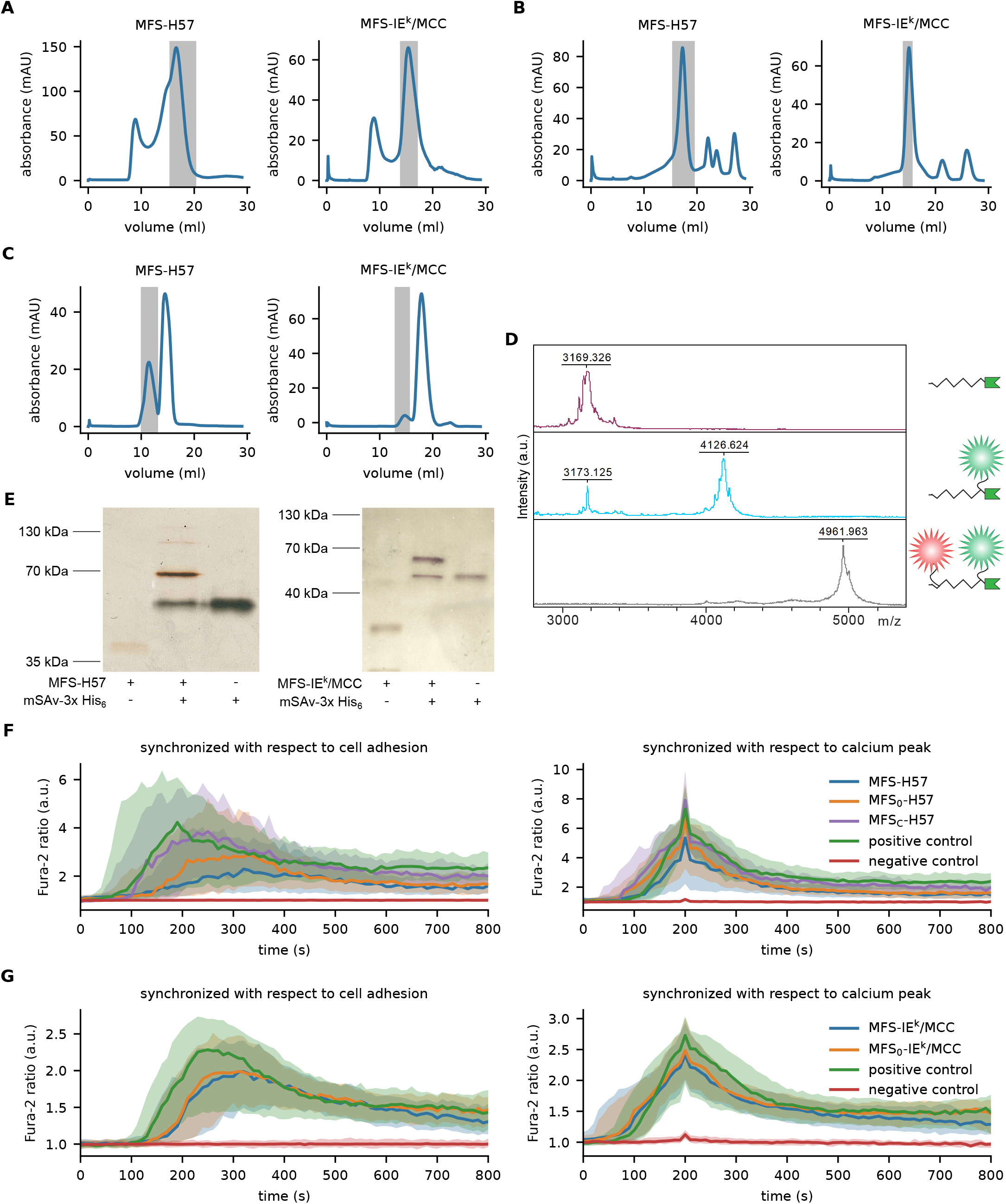
Synthesis and functional validation of the TCR-specific force sensor. (**A**) Refolded single chain antibody fragment derived from the TCRβ-reactive H57 monoclonal antibody (H57-scF_V_) and refolded murine pMHC class II molecule IE^k^ with MCC peptide (IE^k^/MCC) was purified by Superdex 200 10/300 GL (S200) size exclusion chromatography. The gray area corresponds to collected fractions. (**B**) Post conjugation, the surplus of DBCO was removed from monomeric H57-scF_V_-DBCO and IE^k^/MCC-DBCO via Superdex 75 10/300 GL (S75) gel filtration. (**C**) Unconjugated MFS was separated from MFS-H57 or MFS-IE^k^/MCC via S75 or S200 gel filtration, respectively. (**D**) Mass spectra of HPLC purified unlabeled, Alexa Flour 555-labeled and Alexa 555 / Alexa 647 dual-labeled MFS (top to bottom) confirmed successful conjugation. The theoretical molar weight of the unconjugated peptide is 3172 Da. (**E**) SDS-PAGE gel shift analysis of MFS-H57 and MFS-IE^k^/MCC in the presence of mSAv-3xHis_6_ performed under non-boiling and non-reducing conditions testifies to the complete conjugation of the force sensor. In case of MFS-H57, a second slight band corresponding to a double shift and a third barely visible band corresponding to triple shift indicated the presence of a small fraction of H57-scF_V_ conjugated to more than one sensor peptide. (**F-G**) 5c.c7 TCR-transgenic T-cell blasts were brought into contact with fluid-phase SLB decorated with 100-150 molecules µm^-2^ of force sensor bound to mSAv-3xHis_6_ together with His-tagged ICAM-1 and B7-1. Fura-2 emission was measured at 340 and 380 nm excitation. The left panels show the increase of intracellular calcium levels measured by ratiometric Fura-2 analysis for MFS-H57, MFS_0_-H57, and MFS_C_-H57 (**F**), and for MFS-IE^k^/MCC and MFS_0_-IE^k^/MCC (**G**). The right panels show the average calcium traces after synchronization with respect to the single cell calcium peaks. SLBs lacking the force sensors served as negative control (only His-tagged ICAM-1), as positive control we used SLBs containing His-tagged ICAM-1, B7-1, and MCC-loaded IE^k^. Median per time point normalized by median of negative control is shown; interquartile range is shown as shaded area; n cells per sample with 112 ≤ n ≤ 311.

**Supplementary Figure 2:**
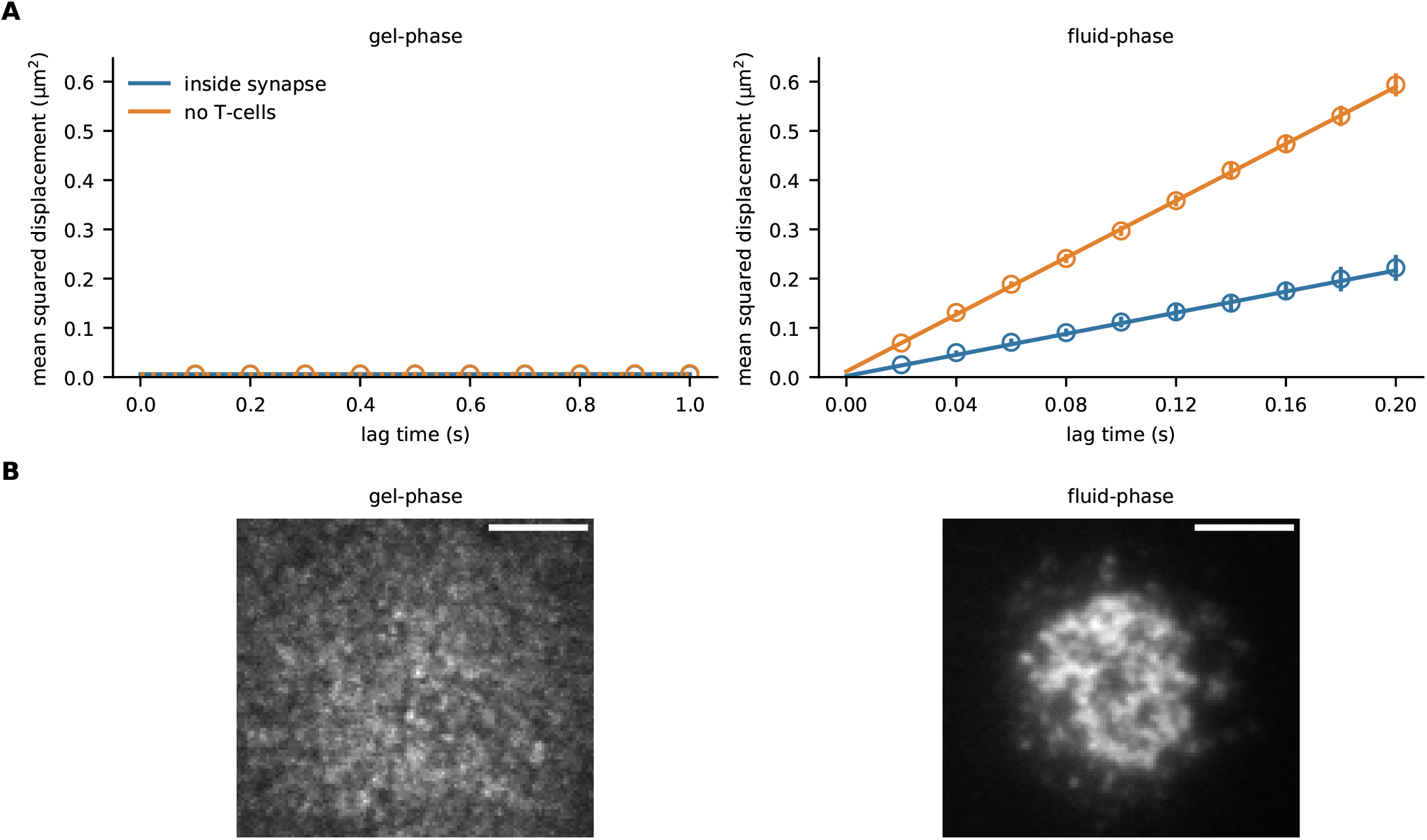
Diffusion analysis of the force sensor anchored to gel-phase and fluid phase SLBs and representative images of the immunological synapses. (**A**) Diffusion analysis of MFS on DPPC membranes yielded *D* = 5.8 ꞏ 10^-5^ ± 8.6 ꞏ 10^-5^ μm² s^-1^ in the presence and *D* = 1.3 ꞏ 10^-4^ ± 6 ꞏ 10^-5^ μm² s^-1^ in the absence of T-cells (left panel). Analogous analysis employing POPC SLBs gave rise to *D* = 0.27 ± 0.02 μm² s^-1^ in the presence and *D* = 0.72 ± 0.02 μm² s^-1^ in the absence of T-cells (right panel). Reduced MFS mobility in the synapse arose from the presence of bound and immobilized molecules (compare Supplementary Fig. 8). 2945 datapoints in 90 tracks from 77 movies for DPPC underneath cells, 3867 datapoints in 102 tracks from 30 movies for DPPC without cells, 724 datapoints in 29 tracks from 109 movies for POPC underneath cells, 904 datapoints in 31 tracks from 30 movies for POPC without cells. (**B**) TIRF imaging of MFS employed at high densities showed microcluster formation within the immunological synapse of T-cells in contact with fluid-phase POPC-based SLBs (right panel) but not of T-cells engaging gel-phase DPPC-based SLBs (left panel). Scale bars 5µm.

**Supplementary Figure 3:**
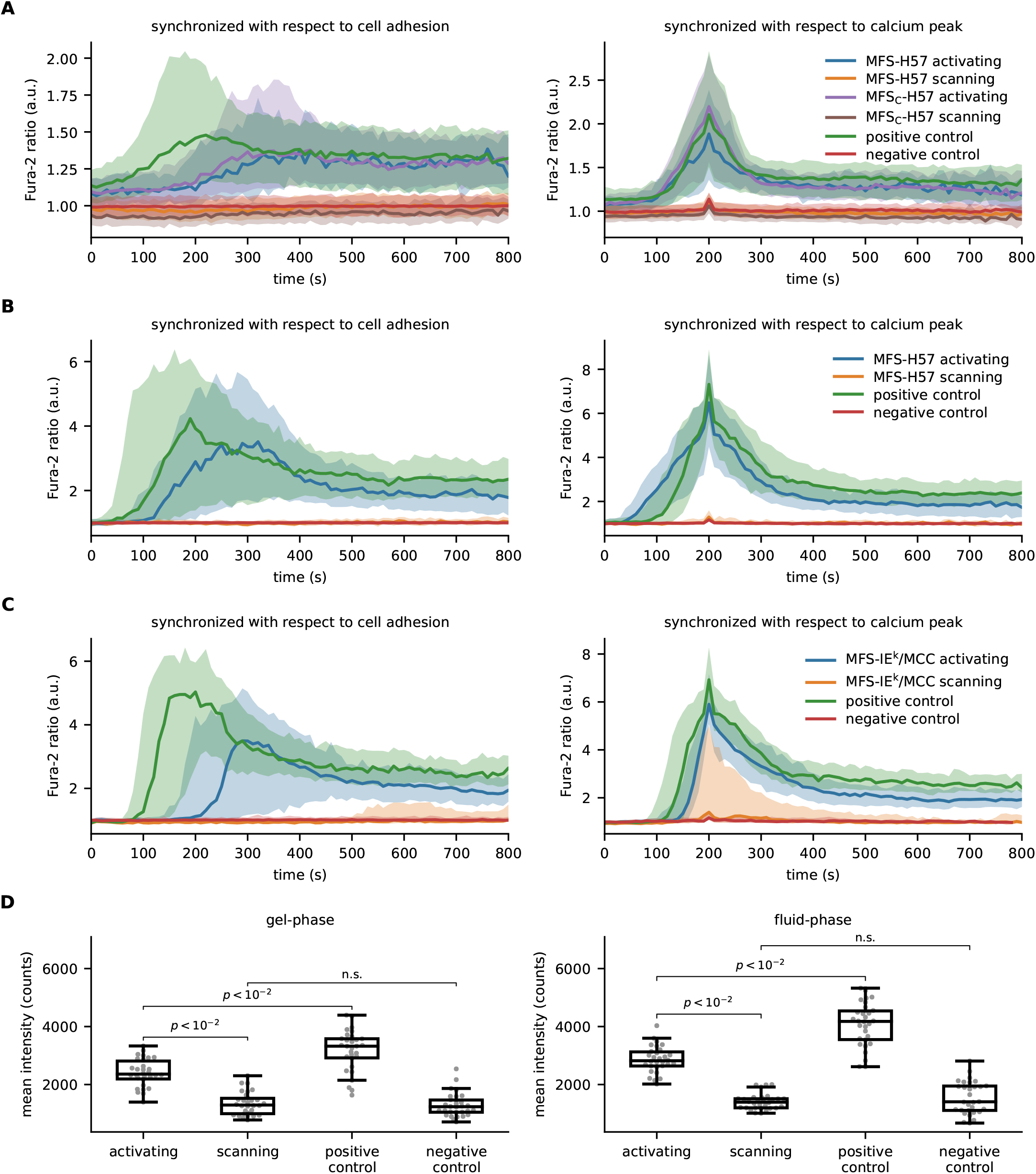
Functional response of T-cells in contact with gel-phase and fluid-phase SLBs displaying the force sensor. (**A-C**) Ratiometric calcium imaging of T-cells confronted with gel-phase DPPC SLBs (**A, C**) and fluid-phase POPC SLBs (**B**). All SLBs employed featured His-tagged ICAM-1. In addition, one of the following (combination of) constructs were added: Panel (**A**): MFS-H57 (single molecule density) + MFS_0_-H57 (high density) + B7-1 (**blue**); MFS-H57 (single molecule density) (**orange**); MFS_C_-H57 (single molecule density) + MFS0-H57 (high density) + B7-1 (**purple**); MFS_C_-H57 (single molecule density) (**brown**). Panel (**B**): MFS-H57 (single molecule density) + MFS_0_-H57 (high density) + B7-1 (**blue**); MFS-H57 (single molecule density) (**orange**). Panel (**C**): MFS-IE^k^/MCC (single molecule density) + MFS_0_-IE^k^/MCC (high density) + B7-1 (**blue**); MFS-IE^k^/MCC (single molecule density) (**orange**). In all panels, SLBs lacking the force sensors served as negative control (only His-tagged ICAM-1; **red**); as positive control we used SLBs containing His-tagged ICAM-1, B7-1, and MCC-loaded IE^k^ (**green**). Median per time point normalized by median of negative control is shown; interquartile range is shown as shaded area. The left panels show the average calcium traces after synchronization with respect to cell adhesion, the right panels after synchronization with respect to peak position. We used between 60 and 367 cells for analysis. (**D**) ZAP70 recruitment to the immunological synapse was assessed via immunostaining in conjunction with brightness analysis and TIRF microscopy. 5c.c7 T-cells were brought in contact with gel-phase (left panel) or fluid-phase SLBs (right panel), under activating or scanning conditions. T-cells were stimulated via MFS-H57. As positive control we used SLBs functionalized with ICAM-1, B7-1, and MCC-loaded IE^k^, as negative control SLBs functionalized only with ICAM-1. Boxes indicate the interquartile range (first quartile to third quartile), whiskers extend 1.5 times the interquartile range from the first and third quartile, the line indicates the median.

**Supplementary Figure 4:**
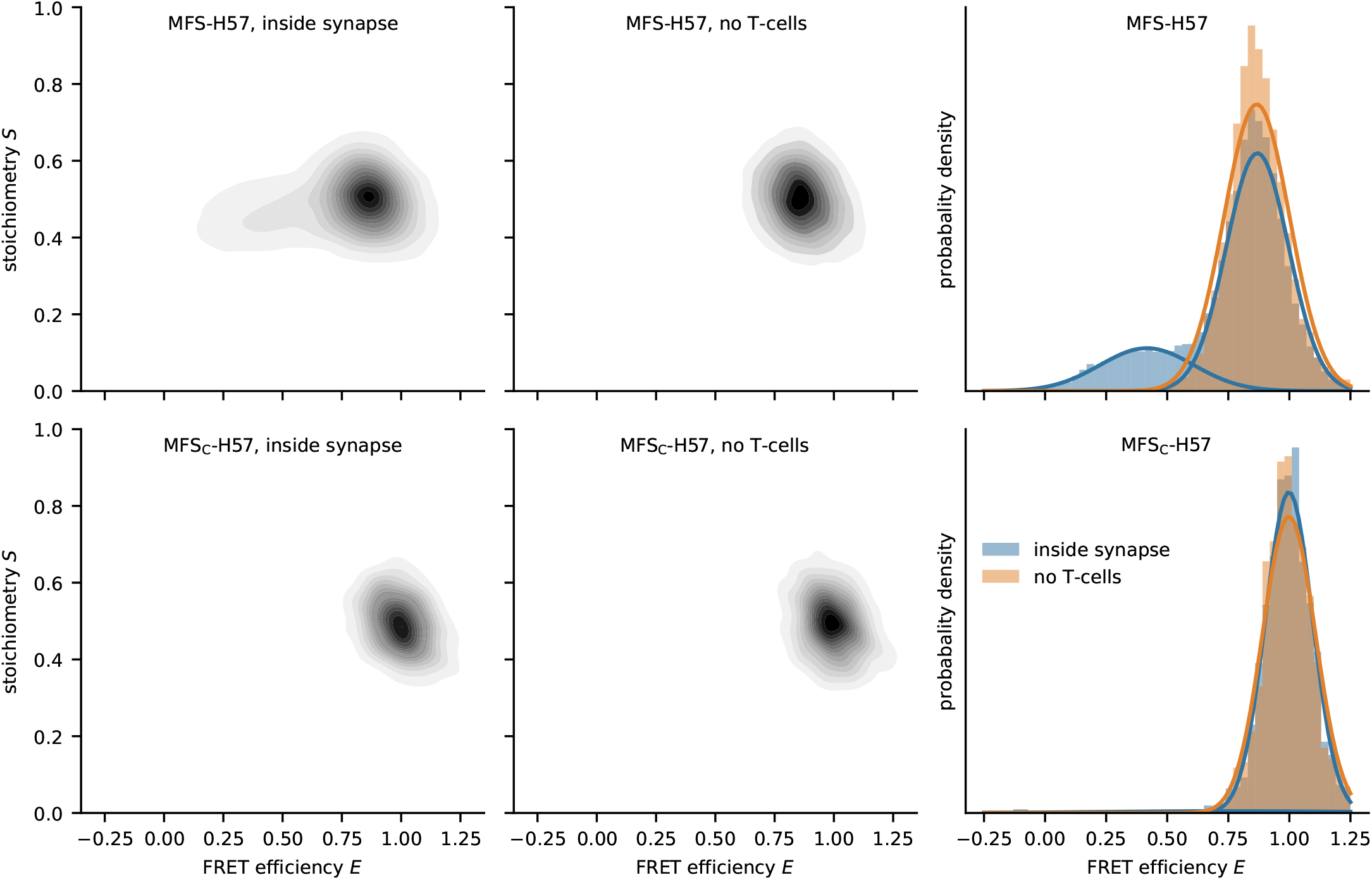
Reading out the force sensor at the single-molecule level. FRET efficiency – stoichiometry (*E-S*) analysis of single-molecule datasets obtained for MFS and MFS_C_ present on gel-phase SLBs in the absence and presence of T-cells. Each data point in the depicted cloud corresponds to one single-molecule FRET event. Measured FRET efficiencies *E* are indicated in histograms (right panels). A Gaussian mixture model was fitted to the *E-S* data, yielding pronounced maxima at *E* = 0.87 and *E* = 0.41 (MFS in the presence of T-cells), *E* = 0.87 (MFS in the absence of T-cells), *E* = 0.99 (MFS_C_ in the presence of T-cells), and E = 1.00 (MFS_C_ in the absence of T-cells). In the presence of T-cells the low FRET peak corresponds to a weight of 32%. n = 15,556 from 593 movies for MFS in the presence of T-cells, n = 12,657 from 195 movies for MFS in the absence of T-cells, n = 1,451 from 127 movies for MFS_C_ in the presence of T-cells, n = 1,506 from 30 movies for MFS_C_ in the absence of T-cells.

**Supplementary Figure 5:**
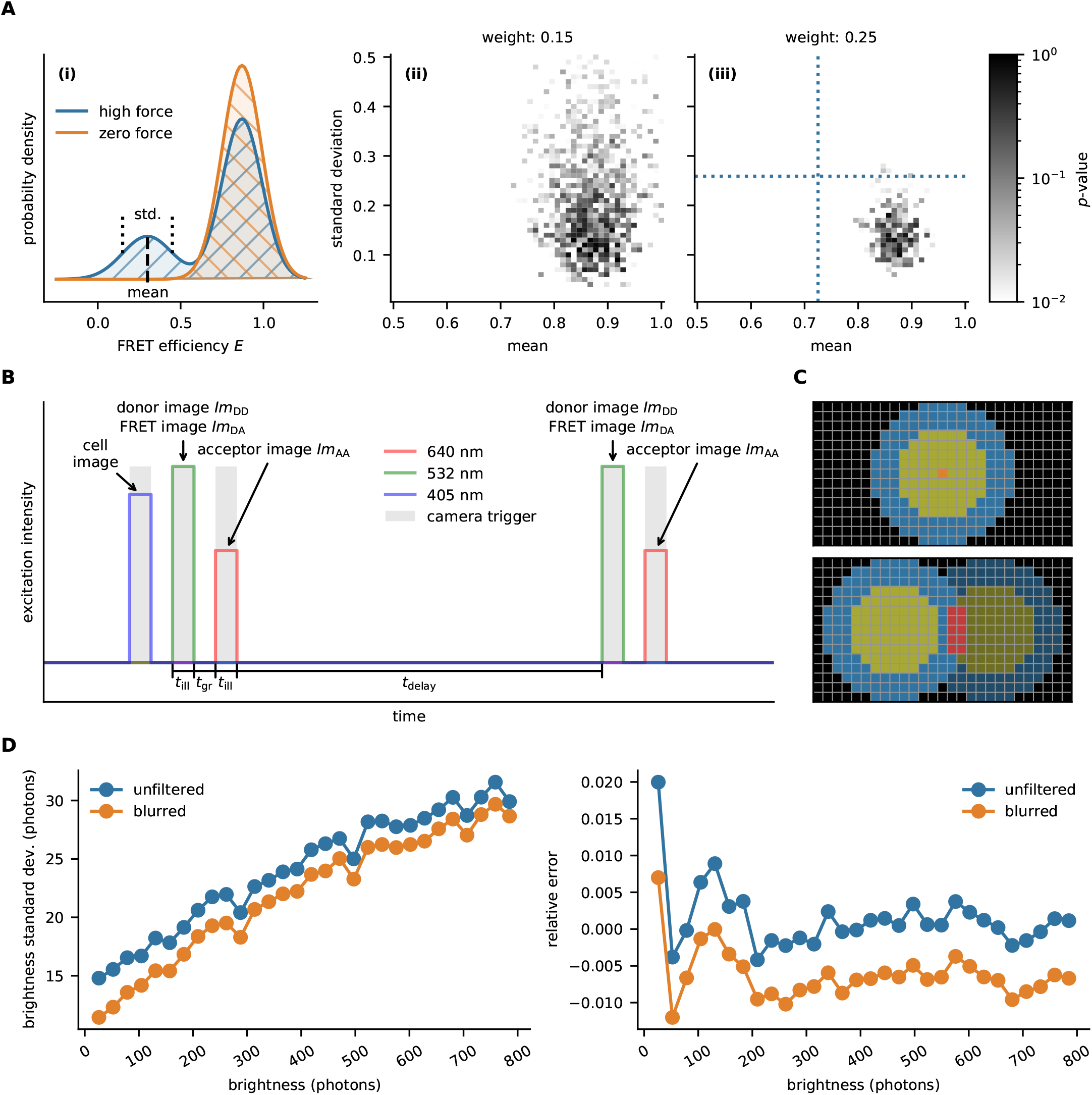
Quantitation of the single-molecule FRET signal. (**A**) Determination of the lower boundary, *F_min_*, of the sensitivity towards applied forces conveyed by the MFS. We simulated Gaussian-distributed random data sets for the high-FRET peak, taking a mean FRET efficiency of 0.87 and a standard deviation *σ* = 0.12. For the low FRET peak, we simulated different values for the mean FRET efficiency and the standard deviation (**i**). Samples were compared to a distribution lacking the low FRET peak using a two-sample Kolmogorov-Smirnov test. Determined p-values were plotted for the different parameter settings. Panel (**ii**) shows results assuming a statistical weight of 0.15 for the low FRET peak, panel (**iii**) for a statistical weight of 0.22. Dashed lines indicate the results from the experiments shown in Supplementary Fig. 7 for MFS-H57 recorded on fluid phase SLBs under scanning conditions on 5c.c7 T cells (mean = 0.73, std = 0.26). (**B**) Illumination timing. Each image sequence is a repetition of two consecutive images recorded with 532 nm and 640 nm excitation, yielding Im_DD_, Im_DA_, and Im_AA_. The timings are indicated in the image. Prior to this, samples were illuminated with 405nm to obtain one image of the outline of cell in the Fura-2 channel. (**C**) Mask used for brightness analysis. If a molecule is located in the orange position, we calculated its brightness by summing over the yellow pixels, and subtracted the weighted blue pixels as background (top). If background of one signal overlaps with the foreground of another signal (bottom image), the corresponding affected pixels (red) were omitted in the analysis. (**D**) Gaussian blur reduces noise in brightness determination (left). For dim signals, noise is decreased by 25 % to 30 %. The blur leads to slight underestimation of the brightness (typically < 1 %, right). Since this is constant over the brightness range, derived ratiometric quantities, like E and S, are not biased.

**Supplementary Figure 6:**
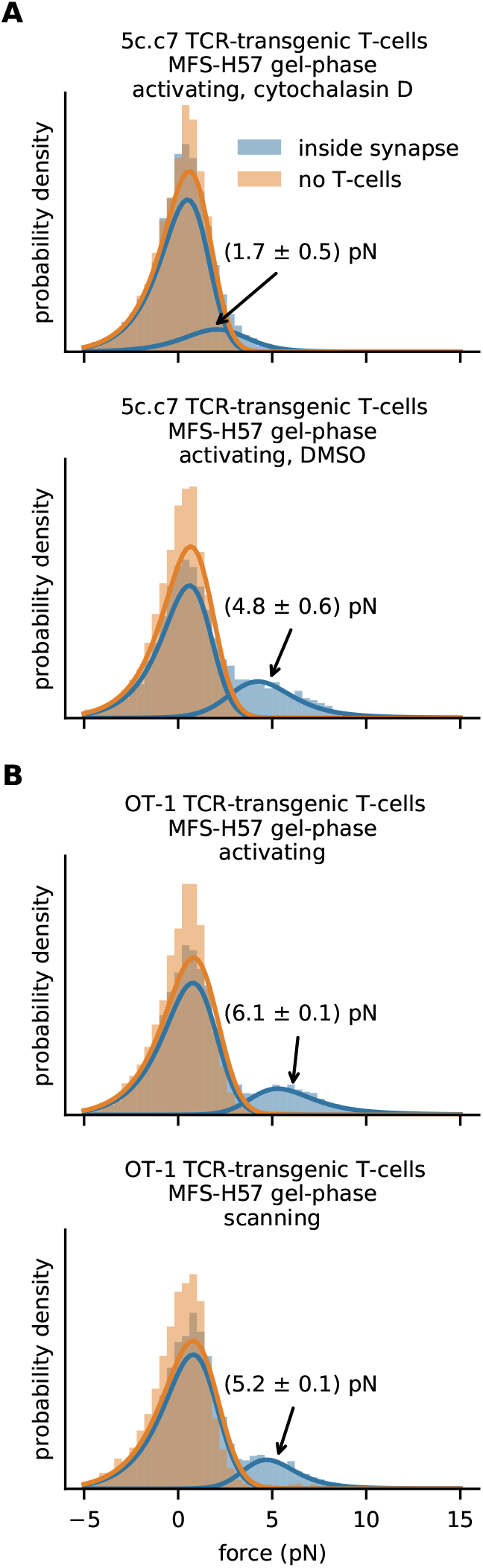
Influence of actin cytoskeleton and T-cell type on TCR-imposed forces. (**A**) Single molecule analysis of MFS-H57 on gel-phase SLBs recorded under activating conditions using 5c.c7 T-cells, upon inhibition of actin polymerization via cytochalasin D (top). The bottom panel shows the vehicle control. We observed a substantial reduction of the high force peak. n = 5,330 (239 movies) for cytochalasin D-treated T-cells, n = 5,390 (70 movies) without T-cells, n = 4,865 (255 movies) for the vehicle control with T-cells, n = 4,637 (69 movies) without T-cells. (**B**) Single molecule analysis of MFS-H57 on gel-phase SLBs recorded under activating (top) or scanning conditions (bottom), using OT-1 CD8^+^ T-cells. We observed a significant high force peak at 6.1 ± 0.1 pN (18%) for the activating case and 5.2 ± 0.1 pN (15%) under scanning conditions. n = 8,963 (408 movies) for activated T-cells, n = 5,450 (119 movies) for stimulatory SLBs without T-cells, n = 5,435 (226 movies) for scanning T-cells, n = 4,572 (66 movies) for non-stimulatory SLBs without T-cells.

**Supplementary Figure 7:**
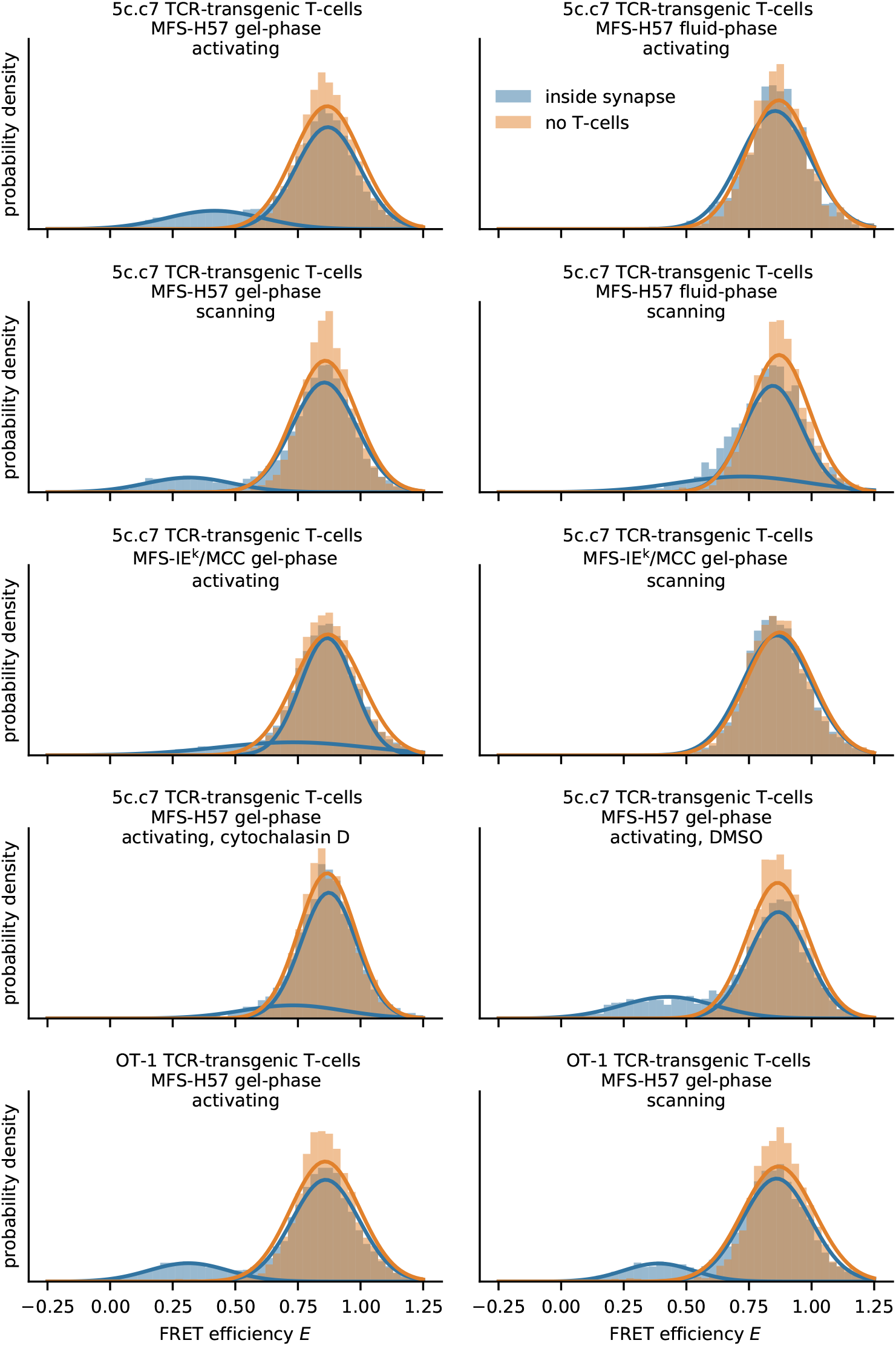
Histograms of single molecule FRET data. Histograms of FRET efficiencies *E* for the data sets shown in Fig. 2, Fig. 4, Supplementary Fig. 6. Lines indicate the fit results of the Gaussian mixture model for SLBs in contact with T-cells, and single Gaussian fits for SLBs without T-cells.

**Supplementary Figure 8:**
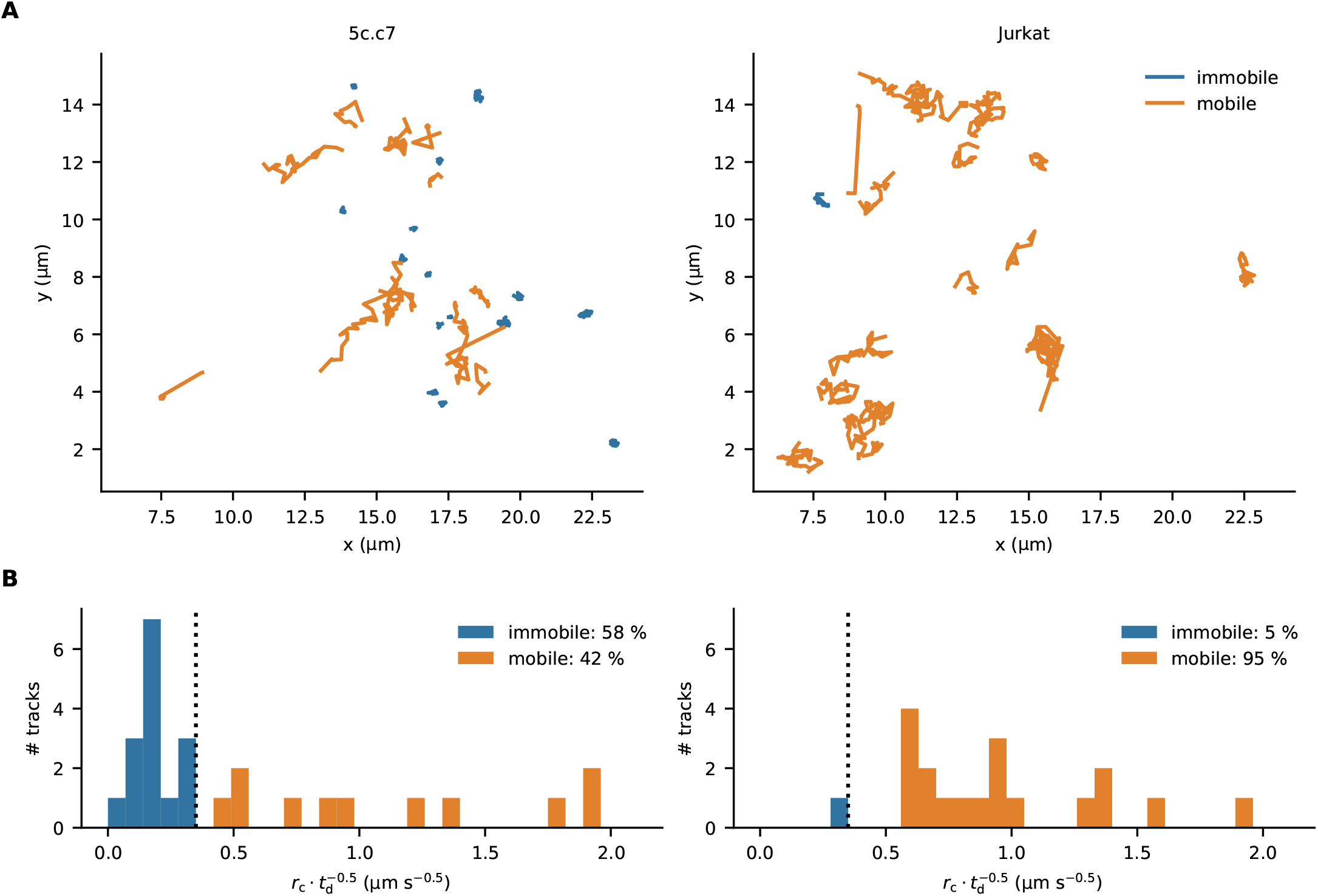
MFS mobility on POPC SLBs is reduced after synaptic TCR binding. Mobility of MFS on POPC SLBs analyzed in a synapse involving 5c.c7 T-cells (left) or Jurkat T-cells serving as a negative control (right). Panel (**A)** shows selected trajectories, panel (**B)** a histogram of the calculated extension of the trajectory, *r*, normalized by the square root of its duration, *t_d_*. The selection threshold is indicated as dashed line. Diffusion analysis of MFS yielded: D=0.018 ± 0.007 μm² s^-1^ and D=0.89 ± 0.17 μm² s^-1^ for the immobile and mobile fraction in a synapse with 5c.c7 T-cells, respectively, and D=0.71 ± 0.1 μm² s^-1^ for the mobile fraction in a synapse with Jurkat T-cells. 724 events in 29 tracks from 109 movies for 5c.c7 cells, 561 events in 20 tracks from 68 movies for Jurkat cells.

**Supplementary Figure 9:**
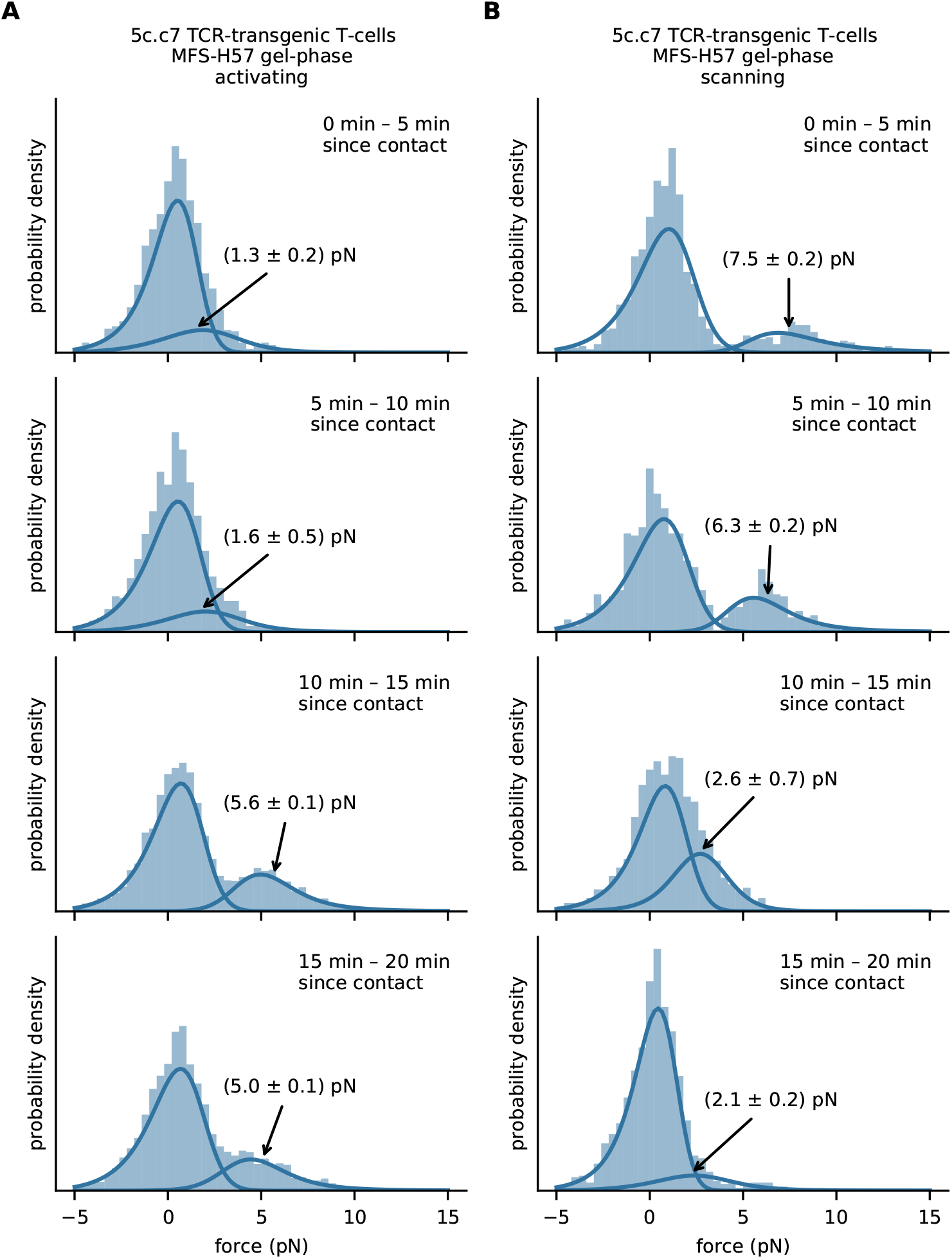
Single molecule forces alter during the time course of T-cell adhesion. We used MFS-H57 data recorded on 5c.c7 T-cells contacting activating (**A**) or non-activating (**B**) gel-phase SLBs. Data were grouped with respect to the time elapsed since the corresponding T-cell first contacted the SLB. Fit results for the average pulling forces are indicated in the figure. The following number of fata points were used. Activating conditions: 0-5 min, n = 1579 (38 movies); 5-10 min, n = 2701 (60 movies); 10 – 15 min, n = 3403 (69 movies); 15 – 20 min, n = 4047 (75 movies). Scanning conditions: 0-5 min, n = 1140 (29 movies); 5 – 10 min, n = 927 (21 movies); 10 – 15 min, n = 1105 (32 movies); 15 – 20 min, n = 1410 (32 movies).

**Supplementary Figure 10:**
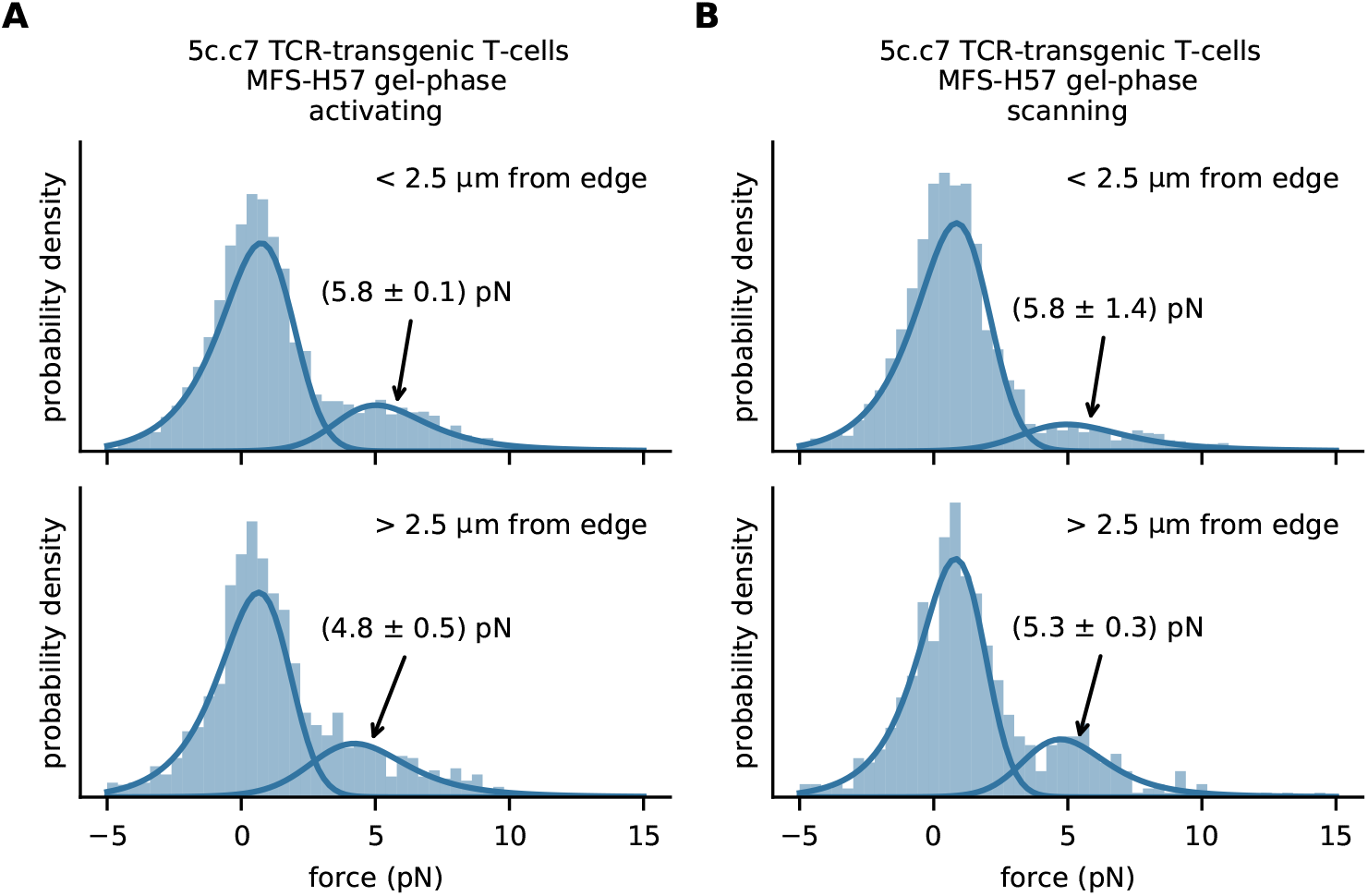
Spatial analysis of single molecule force histograms. We used MFS-H57 data recorded on 5c.c7 T-cells contacting activating (**A**) or non-activating (**B**) gel-phase SLBs. Data were grouped with respect to the closest distance of the FRET event from the cell border, representing the periphery (0 – 2.5 µm) or the center of the T-cell (> 2.5 µm). The Gaussian mixture model revealed the indicated average forces for the high force peak. Activating conditions: < 2.5 μm from edge, n = 4650 (105 movies); > 2.5 μm from edge, n = 1498 (50 movies). Scanning conditions: < 2.5 μm from edge, n = 3367 (92 movies); > 2.5 μm from edge, n = 690 (21 movies).

**Supplementary Figure 11:**
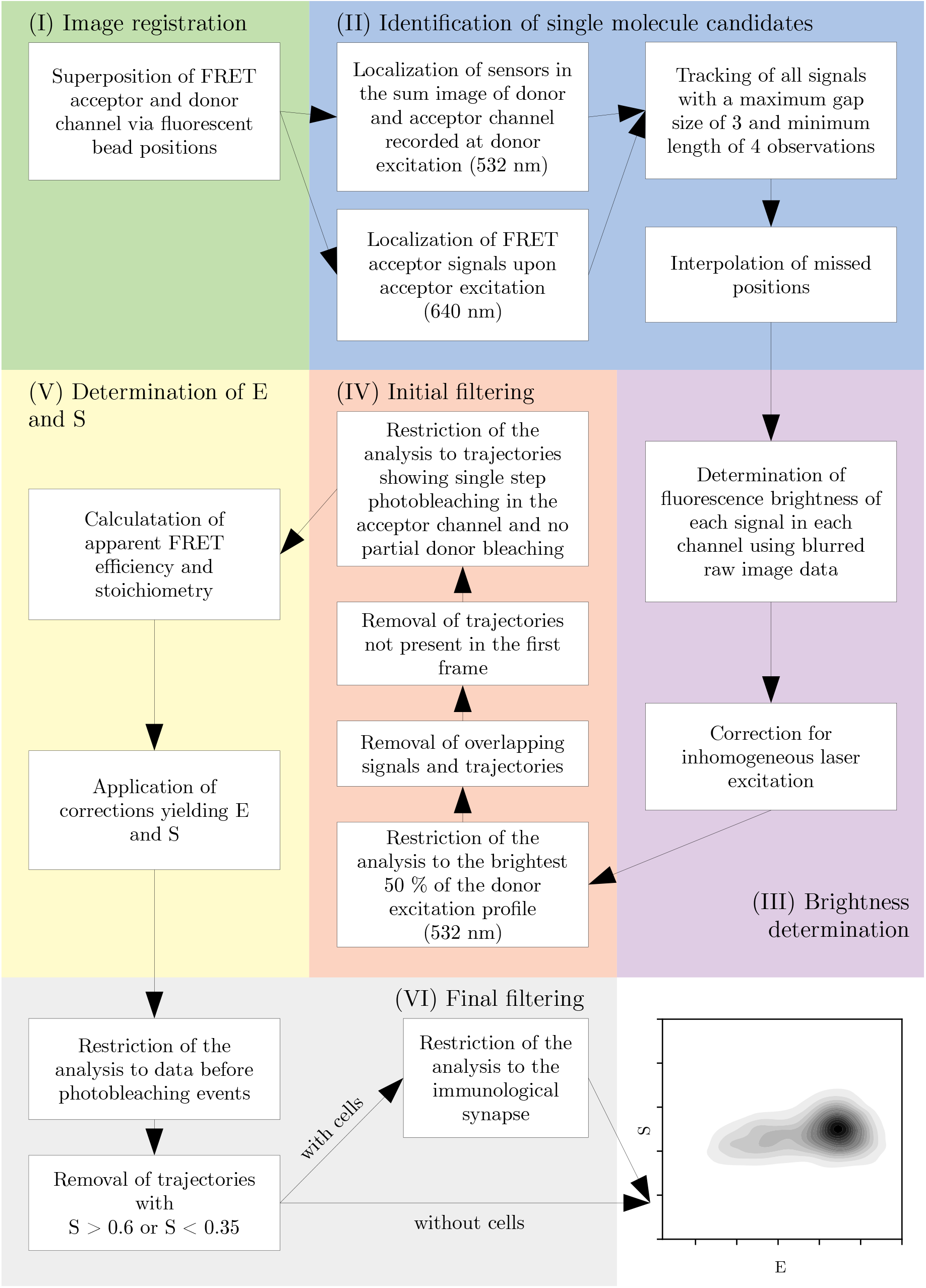
FRET analysis workflow. Flow chart describing the single-molecule FRET analysis workflow. Subheadings relate to the description in the Materials and Methods section.

**Supplementary Table 1:** The results of a fit with a two-component Gaussian mixture model yielded the results shown in the table.

| condition |  | low-FRET component |  |  | high-FRET component |  |  |
| --- | --- | --- | --- | --- | --- | --- | --- |
|  |  | mean | std. | weight | mean | std. | weight |
| 5c.c7 MFS-H57 gel-phase activating | inside synapse | 0.42 | 0.19 | 0.21 | 0.87 | 0.13 | 0.79 |
|  | no T-cells | 0.89 | 0.29 | 0.09 | 0.86 | 0.11 | 0.91 |
| OT-1 MFS-H57 gel-phase activating | inside synapse | 0.31 | 0.16 | 0.18 | 0.86 | 0.13 | 0.82 |
|  | no T-cells | 0.82 | 0.30 | 0.10 | 0.86 | 0.10 | 0.90 |
| 5c.c7 MFS-H57 gel-phase activating, DMSO | inside synapse | 0.43 | 0.19 | 0.24 | 0.87 | 0.12 | 0.76 |
|  | no T-cells | 0.78 | 0.36 | 0.04 | 0.87 | 0.10 | 0.96 |
| 5c.c7 MFS-H57 gel-phase activating, cytochalasin D | inside synapse | 0.73 | 0.20 | 0.16 | 0.87 | 0.11 | 0.84 |
|  | no T-cells | 0.86 | 0.10 | 0.82 | 0.91 | 0.18 | 0.18 |
| 5c.c7 MFS-H57 gel-phase scanning | inside synapse | 0.32 | 0.17 | 0.15 | 0.86 | 0.13 | 0.85 |
|  | no T-cells | 0.74 | 0.31 | 0.06 | 0.87 | 0.10 | 0.94 |
| OT-1 MFS-H57 gel-phase scanning | inside synapse | 0.39 | 0.14 | 0.15 | 0.86 | 0.14 | 0.85 |
|  | no T-cells | 0.79 | 0.34 | 0.08 | 0.88 | 0.11 | 0.92 |
| 5c.c7 MFS-IE<sup>k</sup>/MCC gel-phase activating | inside synapse | 0.73 | 0.30 | 0.24 | 0.87 | 0.11 | 0.76 |
|  | no T-cells | 0.86 | 0.11 | 0.90 | 0.94 | 0.27 | 0.10 |
| 5c.c7 MFS-IE<sup>k</sup>/MCC gel-phase scanning | inside synapse | 0.84 | 0.12 | 0.77 | 0.90 | 0.23 | 0.23 |
|  | no T-cells | 0.85 | 0.11 | 0.82 | 0.98 | 0.19 | 0.18 |
| 5c.c7 MFS-H57 fluid-phase activating | inside synapse | 0.79 | 0.30 | 0.10 | 0.86 | 0.12 | 0.90 |
|  | no T-cells | 0.85 | 0.10 | 0.75 | 0.92 | 0.21 | 0.25 |
| 5c.c7 MFS-H57 fluid-phase scanning | inside synapse | 0.73 | 0.26 | 0.25 | 0.85 | 0.12 | 0.75 |
|  | no T-cells | 0.86 | 0.09 | 0.75 | 0.90 | 0.19 | 0.25 |

